# Tractometry reproducibility and generalizability across scanners, scanner models, and acquisition protocols

**DOI:** 10.64898/2026.05.13.723388

**Authors:** Daiki Taguma, Isao Yokoi, Takuji Kinjo, Shuhei Tsuchida, Toshikazu Miyata, Tetsuya Matsuda, Garikoitz Lerma-Usabiaga, Hiromasa Takemura

## Abstract

Diffusion-weighted magnetic resonance imaging (dMRI)-based tractometry enables the quantification of white matter tissue properties in living humans while preserving anatomical specificity. Although tractometry is highly reproducible when the same scanner and acquisition protocol are used, its generalizability across scanners and protocols remains unclear. To address this gap, we performed a traveling-head experiment involving five subjects to evaluate tractometry across progressively different acquisition conditions, including multiple scanners, different scanner models, and two distinct protocols. Tractometry was performed for 20 major white matter tracts using diffusion tensor imaging metrics, neurite orientation dispersion and density imaging (NODDI) metrics, and a semi-quantitative ratio metric (T1w/b0). Generalizability across dataset pairs was quantified using the intraclass correlation coefficient (ICC). Tractometry showed consistently high ICCs when the scanner and protocol were identical; however, ICCs declined as differences in scanner model and acquisition protocol increased. Fractional anisotropy and orientation dispersion index retained relatively high ICCs across these comparisons, whereas other metrics showed marked declines when scanners or protocols differed. ComBat harmonization partially mitigated these declines, but ICCs did not reach the levels observed for datasets acquired using identical scanners and protocols. Finally, the minimum detectable change (MDC) for tractometry in datasets pooled across scanners and protocols varied by tract; for example, the optic radiation showed a lower MDC than the cingulum hippocampus. These findings highlight both the strengths and limitations of tractometry in multisite studies and highlight the importance of quantifying scanner- and protocol-dependent effects for specific metrics and tracts when interpreting measurements from heterogeneous datasets.

## 1. Introduction

Quantifying human white matter tissue properties in health and disease is central to neuroscience research on disorders and the neural basis of cognitive functions (Forkel et al., 2022; Takemura et al., 2024; Thomason & Thompson, 2011). Diffusion-weighted magnetic resonance imaging (dMRI) is an essential tool for this purpose because it enables non-invasive measurement of white matter tissue properties in living humans, thereby supporting investigations across a wide range of clinical and neuroscientific questions (Assaf et al., 2019; Mori & Zhang, 2006; Wandell, 2016). Several strategies are available for analyzing human dMRI data, including region-of-interest (ROI) analyses, volume and shape analyses of white matter tracts, and network analyses of structural connectivity (Amemiya et al., 2021; Glozman et al., 2018; Lebel et al., 2019; Sotiropoulos & Zalesky, 2019).

Among these approaches, tractometry identifies major white matter tracts using tractography and then evaluates tissue properties along each tract trajectory based on dMRI metrics, such as mean diffusivity (MD) (Gerig et al., 2004; Jones et al., 2005; Wakana et al., 2007; Yeatman et al., 2012; Yendiki et al., 2011). A key advantage of tractometry is that it constrains the analysis to white matter tracts known to exist in anatomical literature, thereby reducing the risk of generating and interpreting false-positive pathways (Maier-Hein et al., 2017; Schilling et al., 2020). Because tractometry supports interpretation based on established white matter anatomy, it also facilitates integration with findings from lesion studies (Catani & Ffytche, 2005) and intervention studies (Alagapan et al., 2023). Several open-source tractometry tools are now available to the research community (Chandio et al., 2020; Kruper et al., 2021; Lerma-Usabiaga et al., 2023; Yendiki et al., 2011), contributing to the widespread use of this approach in basic and clinical neuroscience.

dMRI-based tractometry generally yields highly reproducible results when data are acquired using the same MRI scanner and identical acquisition protocol (Cousineau et al., 2017; Kruper et al., 2021; Lerma-Usabiaga et al., 2023; Liu et al., 2022). However, dMRI-based metrics such as MD are sensitive to differences in acquisition parameters, including b-values, and MRI hardware (Grech-Sollars et al., 2015; Lerma-Usabiaga et al., 2019; Saito et al., 2023). This variability complicates comparisons across research groups and limits the ability to draw generalizable conclusions from heterogeneous populations and multinational cohorts. Therefore, rigorous evaluation of the dependence of tractometry on scanner models and acquisition protocols is essential for improving its generalizability. Hereafter, we define “reproducibility” as the consistency of tractometry data obtained using the same scanner and protocol, whereas “generalizability” refers to the consistency of results across different scanners and protocols.

Here, we evaluated the reproducibility and generalizability of tractometry to complement findings from previous multisite dMRI studies (Tong et al., 2019, 2020; Vollmar et al., 2010). We used a traveling-head design (Jovicich et al., 2006; Tong et al., 2020; Yamashita et al., 2019) to acquire data from the same subjects across multiple scanners and acquisition protocols. Specifically, the experimental design enabled comparisons across progressively different acquisition conditions: within the same scanner, between identical scanner models, and across different scanner models. This design allowed us to estimate the degree of inconsistency in tractometry results attributable to these differences. We acquired datasets using 3T MRI scanners from two models, Verio and PrismaFit (Figure 1). Five subjects participated in two sessions on separate dates for each scanner, and two acquisition protocols were tested to evaluate protocol-dependent effects. Data were processed using an established tractometry pipeline, RTP2 (Lerma-Usabiaga et al., 2023), to extract diffusion tensor imaging (DTI) metrics, neurite orientation dispersion and density imaging (NODDI) metrics (Zhang et al., 2012), and a semi-quantitative T1w/b0 ratio metric (Moskovich et al., 2024). Tractometry reproducibility and generalizability were quantified using the intraclass correlation coefficient (ICC), which assessed agreement across dataset pairs. We also evaluated whether ComBat harmonization improved generalizability (Fortin et al., 2017, 2018; Johnson et al., 2007). Finally, we quantified the minimum detectable change (MDC) of tractometry in datasets pooled across scanners and protocols for individual tracts to identify tracts that were relatively robust to heterogeneous acquisition conditions. Overall, this study provides insight into how hierarchical differences in hardware and protocols affect tractometry, the extent to which harmonization mitigates these dependencies, and the sensitivity of specific metrics and tracts to scanner- and protocol-related factors.

**Figure 1.**
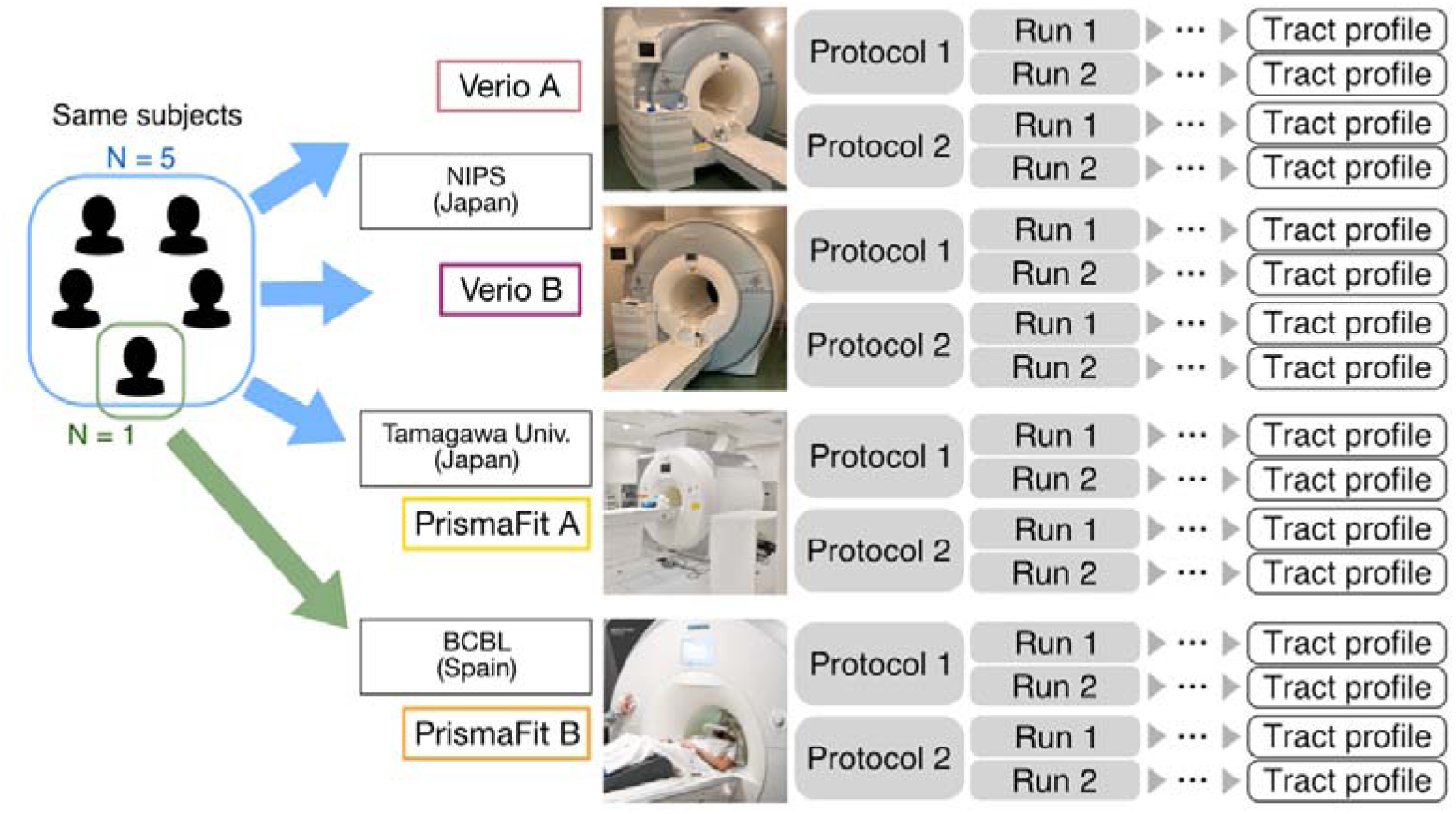
Experimental design. A traveling-head experiment was conducted in five subjects using three 3T MRI scanners in Japan. One subject also participated in the experiment at the Basque Center on Cognition, Brain and Language (BCBL), Spain. In all experiments, two dMRI acquisition protocols were tested. Each dMRI acquisition was repeated on a separate date to evaluate test–retest reproducibility.

## 2. Methods

### 2.1. Subjects

Five healthy male volunteers (ages 24, 28, 32, 36, and 48 years; mean age, 34.0 years) traveled to each site for scanning with three scanners: two Siemens 3T Verio scanners at the National Institute for Physiological Sciences (NIPS), Japan, and one Siemens 3T PrismaFit scanner at Tamagawa University, Japan. One volunteer (age 25 years; subject 2) also participated in the experiment at the Basque Center on Cognition, Brain, and Language (BCBL), Spain. The experimental protocol was approved by the Ethics Committee of the National Institutes of Natural Sciences (protocol number: EC01-093), the Ethics Committee of Tamagawa University (protocol number: TRE23-0002), and the Ethics Committee of the BCBL (protocol number: 190924SM). All subjects provided written informed consent after receiving an explanation of the voluntary nature of participation, MRI safety procedures, the right to withdraw, and the sharing of anonymized data.

The design of traveling-head experiments involves a tradeoff. One approach is to recruit many subjects for scanning with two scanners. This design is effective for evaluating group-level measurements but does not assess how tractometry varies within the same subject across multiple instruments and acquisition protocols. Another approach is to recruit a smaller number of subjects for repeated measurements across more than two scanners, multiple institutions, and multiple acquisition protocols. Although this design is less suitable for group-level evaluation, it provides important insight into within-subject variability and the generalizability of tractometry across scanners and protocols. Similar to previous studies that included small numbers of traveling subjects (N = 1, Palacios et al., 2017; N = 3, Tong et al., 2019), we used the latter design to evaluate within-subject reproducibility, scanner and protocol dependence, and tractometry generalizability. The number of subjects recruited in this study (N = 5) is comparable to or larger than that in previous traveling-subject studies using a similar design and is appropriate for our objective.

### 2.2. Data acquisition methods

#### 2.2.1. Instrument

This experiment used four 3T MRI systems (Figure 1). First, two Siemens MAGNETOM Verio scanners (Siemens Healthcare GmbH, Erlangen, Germany) at NIPS, Okazaki, Japan, were used with 32-channel head coils. These two scanners, referred to as Verio A and Verio B, were installed in the same year (2009) and had the same maximum gradient strength (40 mT/m). Second, a Siemens MAGNETOM PrismaFit scanner, referred to as PrismaFit A (Siemens Healthcare GmbH), at the Tamagawa University Brain Science Institute, Machida, Japan, was used with a 32-channel head coil. Third, a Siemens MAGNETOM PrismaFit scanner, referred to as PrismaFit B, at the BCBL, Donostia-San Sebastián, Spain, was used with a 64-channel head/neck coil. The PrismaFit scanners had a higher maximum gradient strength than the Verio scanners (80 vs. 40 mT/m).

All experiments using Verio A and Verio B were conducted over two consecutive days. The PrismaFit A experiment at Tamagawa University was performed 1 month after the NIPS experiments using Verio A and Verio B. Given this short interval, long-term white matter alterations, such as aging-related changes, were unlikely to have substantially contributed to variability across these instruments. However, the PrismaFit B experiment at BCBL was performed 10 months after the NIPS experiments.

#### 2.2.2. T1-weighted image acquisition

In each experiment, whole-brain T1-weighted (T1w) images were acquired using magnetization-prepared rapid gradient echo (MP-RAGE) with the following parameters: voxel size, 1 mm isotropic; field of view (FOV), 256 × 240 × 224; repetition time (TR), 2400 ms; echo time (TE), 1.98 ms; flip angle, 8°; inversion time, 1060 ms; iPAT, 2; averages, 2; acquisition time, 5 min 26 s. These images were used for white matter segmentation and ROI identification.

#### 2.2.3. dMRI image acquisition

Two dMRI protocols were used in this study (Table 1). Each was selected based on widely used protocols: protocol 1 was based on the UK Biobank protocol (Miller et al., 2016), and protocol 2 was based on the HARP protocol (Koike et al., 2021). Acquisition parameters were kept identical across all scanners, except for TE, for which the minimum achievable value for each scanner was used. In each run, low b-value images (b = 0 s/mm²) were acquired with reversed phase encoding direction (anterior–posterior) to correct susceptibility-induced geometric distortion. Each measurement was repeated twice to assess within-scanner reproducibility (Figure 1).

**Table 1.** Acquisition parameters for dMRI data. FoV: field of view, dir: directions, iPAT: integrated Parallel Acquisition Techniques, MB: multi-band, TE: echo time, TR: repetition time, PA: posterior-anterior, PE: phase encoding, PF: partial Fourier.

| | FoV<br>(mm) | Voxel<br>Size<br>(mm) | PE<br>dir | MB<br>factor | iPAT | PF | TR<br>(ms) | TE<br>(ms) | b<br>($\text{s/mm}^2$) | # of<br>dir | Scan<br>time<br>(min) |
| --- | --- | --- | --- | --- | --- | --- | --- | --- | --- | --- | --- |
| Protocol 1 | 210×<br>210×<br>144 | 2 iso | PA | 3 | None | 6/8 | 3600 | 90.8<br>(Verio)<br>72.8 | 0/1000/<br>2000 | 6/50/<br>50 | 7:05<br>(Verio)<br>7:08 |
|  |  |  |  |  |  |  |  | (Prisma<br>Fit) |  |  | (Prisma<br>Fit) |
| Protocol 2 | 200×<br>200×<br>148 | 1.7<br>iso | PA | 3 | 2 | 6/8 | 4000 | 90.0<br>(Verio)<br>62.0<br>(Prisma<br>Fit) | 0/700/<br>2000 | 6/20/<br>40 | 6:08<br>(Verio)<br>6:52<br>(Prisma<br>Fit) |

### 2.3. Data preprocessing and tractometry methods

The RTP2 pipeline (Lerma-Usabiaga et al., 2023) was used to process the dMRI and T1w datasets. RTP2 is a containerized pipeline that supports computational reproducibility and data provenance through the launchcontainer Python package (https://github.com/garikoitz/launchcontainers). The RTP2 pipeline organizes the analysis workflow into four main stages: standardizing file formats and folder structures, defining ROIs for tractography, preprocessing dMRI data, and identifying specific tracts for tractometry. In the fourth stage, tractography is performed using the preprocessed dMRI data from the third stage and the ROIs generated in the second stage. The steps implemented in the RTP2 pipeline are briefly described below.

#### 2.3.1. Standardization of file format and folder structure

In the first stage, DICOM data exported from the MRI scanners were converted to NIfTI format and organized according to the Brain Imaging Data Structure (Gorgolewski et al., 2016). Conversion and organization were performed using HeuDiConv (v1.1.0; Halchenko et al., 2025; https://github.com/nipy/heudiconv) and dcm2niix (v1.0.20220720; Li et al., 2016; https://github.com/rordenlab/dcm2niix).

#### 2.3.2. T1w image processing and ROI generation

The second stage of the RTP2 pipeline, RTP2-Freesurferator (v0.2.0_7.4.1rc21; referred to as RTP2-Freesurferator and RTP2-anatROIs in the original paper), processed each subject’s structural T1w images acquired on Verio A to identify ROIs for tract identification. This stage used each subject’s T1w image and ROIs defined in MNI space as inputs and generated a segmented T1w image and subject-specific ROIs aligned to the native T1w space. In all subsequent analyses, each dMRI dataset was co-registered with the T1w image acquired on Verio A during the first run to maintain consistent ROI definition across all dMRI datasets.

The ROIs required for identifying each fiber tract were prepared as follows. First, FreeSurfer (http://surfer.nmr.mgh.harvard.edu/; Fischl, 2012) was used for cortical and subcortical segmentation and parcellation. Thalamic nuclei were then delineated using the FreeSurfer thalamic segmentation module, which relies on a probabilistic atlas constructed from histological and high-resolution *ex vivo* MRI data (Iglesias et al., 2018). In this study, only the lateral geniculate nucleus (LGN) was selected as an ROI. To parcellate the early visual cortex, the Benson atlas (Benson et al., 2012, 2014), implemented in neuropythy (https://github.com/noahbenson/neuropythy), was applied to the FreeSurfer outputs. For optic radiation (OR) identification, V1 and V2 ROIs derived from the Benson atlas were combined to create the visual cortex ROI because macaque tracer studies have shown OR projections to V2 (Yukie and Iwai 1981; Briggs et al. 2016; Liu et al. 2022), and including V2 increases sensitivity for OR identification.

All other ROIs were obtained from atlases in MNI space (Wakana et al., 2007; https://github.com/vistalab/vistasoft/tree/master/mrDiffusion/templates/MNI_JHU_tracts_ROIs; Table 2), following protocols used in previous studies (Yeatman et al., 2012; Zhang et al., 2008). To transform these ROIs into each subject’s native space, Advanced Normalization Tools (ANTs; http://stnava.github.io/ANTs/) were used for nonlinear registration to an MNI template with 1 mm isotropic voxels. Each ROI was expanded by one cubic voxel to ensure that it reached the gray–white matter boundary.

**Table 2.** White matter tracts and ROI files used for tract identification. The table lists the white matter tracts identified from whole-brain streamlines and the corresponding ROI files used for tract identification. ROI files were adapted from Wakana et al. (2007) and are identical to those available as a part of the MATLAB AFQ toolbox (Yeatman et al., 2012).

| Tract | ROI1 | ROI2 | ROI3 |
| --- | --- | --- | --- |
| Left_Thalamic_Radiation | ATR_roi1_L_subj | ATR_roi2_L_subj |  |
| Right_Thalamic_Radiation | ATR_roi1_R_subj | ATR_roi2_R_subj |  |
| Left_Corticospinal | CST_roi1_L_subj | CST_roi2_L_subj |  |
| Right_Corticospinal | CST_roi1_R_subj | CST_roi2_R_subj |  |
| Left_Cingulum_Cingulate | CGC_roi1_L_subj | CGC_roi2_L_subj |  |
| Right_Cingulum_Cingulate | CGC_roi1_R_subj | CGC_roi2_R_subj |  |
| Left_Cingulum_Hippocampus | HCC_roi1_L_subj | HCC_roi2_L_subj |  |
| Right_Cingulum_Hippocampus | HCC_roi1_R_subj | HCC_roi2_R_subj |  |
| Callosum_Forceps_Major | FP_R_subj | FP_L_subj |  |
| Callosum_Forceps_Minor | FA_L_subj | FA_R_subj |  |
| Left_ILF | ILF_roi1_L_subj | ILF_roi2_L_subj | ILF_roi3_L_subj |
| Right_ILF | ILF_roi1_R_subj | ILF_roi2_R_subj | ILF_roi3_R_subj |
| Left_SLF | SLF_roi1_L_subj | SLF_roi2_L_subj |  |
| Right_SLF | SLF_roi1_R_subj | SLF_roi2_R_subj |  |
| Left_Uncinate | UNC_roi1_L_subj | UNC_roi2_L_subj |  |
| Right_Uncinate | UNC_roi1_R_subj | UNC_roi2_R_subj |  |
| Left_Arcuate | SLF_roi1_L_subj | SLFt_roi2_L_subj |  |
| Right_Arcuate | SLF_roi1_R_subj | SLFt_roi2_R_subj |  |

#### 2.3.3. dMRI data preprocessing

The third stage of the RTP2 pipeline, RTP2-preproc (v0.2.4_3.0.4rc1), preprocesses the dMRI data and aligns them to the anatomical reference. This stage follows the established preprocessing protocol implemented in MRtrix3 (Tournier et al., 2019), in combination with ANTs and FSL (Jenkinson et al., 2012). Several MRtrix3 functions were applied sequentially. First, the data were denoised using random matrix theory to exploit redundancy in local principal component analysis (Cordero-Grande et al., 2019; Veraart et al., 2016) through the *dwidenoise* tool. Second, Gibbs ringing artifacts were corrected using *mrdegibbs* (Kellner et al., 2016). Third, susceptibility-induced distortions, eddy-current distortions, and head motion were corrected using FSL’s topup and eddy modules (Andersson et al., 2003; Andersson & Sotiropoulos, 2016), executed through *dwifslpreproc*. Fourth, B_1_ bias field inhomogeneity was corrected using *dwibiascorrect*, and Rician background noise was reduced using *mrcalc*. Finally, a rigid-body transformation was computed using ANTs to register the dMRI data to each subject’s T1w structural image.

#### 2.3.4. Tractography and tractometry

In the fourth and final main stage, the RTP2 pipeline container (v0.2.2_3.0.4rc1) was used to identify white matter tracts from the dMRI data and calculate tract profiles.

*Generation of the whole-brain tractogram:* Fiber orientation distributions (FODs) were estimated for all white matter voxels from the preprocessed dMRI data using MRtrix3’s multi-tissue constrained spherical deconvolution (CSD; Jeurissen et al., 2014), which resolves crossing fibers within individual voxels. CSD-based probabilistic tractography was then performed using the iFOD2 algorithm in MRtrix3 (Tournier et al., 2010) to generate streamlines. To reduce dependence on tractography parameter selection, we used an ensemble tractography approach (Takemura et al., 2016). Specifically, 16 million streamlines were generated by combining eight parameter sets: angle thresholds of 45°, 45°, 45°, 25°, 25°, 25°, 10°, and 10° and maximum streamline lengths of 100, 150, 200, 100, 150, 200, 150, and 200 mm, respectively. For each parameter set, 2 million streamlines were generated using the whole-brain white matter mask as the seed region. The FOD amplitude stopping criterion was fixed at 0.05 across all parameter settings. The streamlines from all parameter sets were concatenated to generate a whole-brain candidate tractogram. The tractogram was then reduced to 500,000 streamlines by excluding streamlines that did not explain the diffusion signal using SIFT (Smith et al., 2013). The remaining streamlines, referred to as the optimized tractogram, were used for subsequent tract identification.

*Optic radiation tractography:* To identify the optic radiation, for which tract endpoints are well established (Sherbondy et al., 2008), the LGN and visual cortical areas V1 and V2 were defined as seed ROIs. A white matter waypoint ROI implemented in RTP2 (OR_roi3_subj) was also used as an inclusion mask. Bidirectional tracking was performed to account for volume differences between the two seed ROIs. To increase tracking sensitivity, the LGN and visual cortex ROIs were dilated into white matter before tracking. For each hemisphere, CSD-based probabilistic tractography using the iFOD2 algorithm in MRtrix3 (Tournier et al., 2010) was used to generate 5,000 streamlines connecting the LGN and visual cortex ROIs. Tracking was constrained by a maximum streamline length of 200 mm, a minimum streamline length of 10 mm, an angle threshold of 45°, and an FOD amplitude stopping criterion of 0.05.

*Tract identification:* Using the RTP2 pipeline container, the tracts of interest (Table 2) were identified from the preprocessed dMRI data by applying AND/NOT operations to streamlines using the ROIs defined in the previous steps (Catani et al., 2002; Mori & van Zijl, 2002). After streamlines belonging to the tracts of interest were selected, outlier streamlines were excluded if their length or spatial position deviated by more than three standard deviations from the mean (Yeatman et al., 2012). For tracts identified from whole-brain streamlines, this procedure was repeated three times. In total, 20 major white matter tracts were identified from all datasets (Table 2), with representative examples shown in Figure 2.

**Figure 2.**
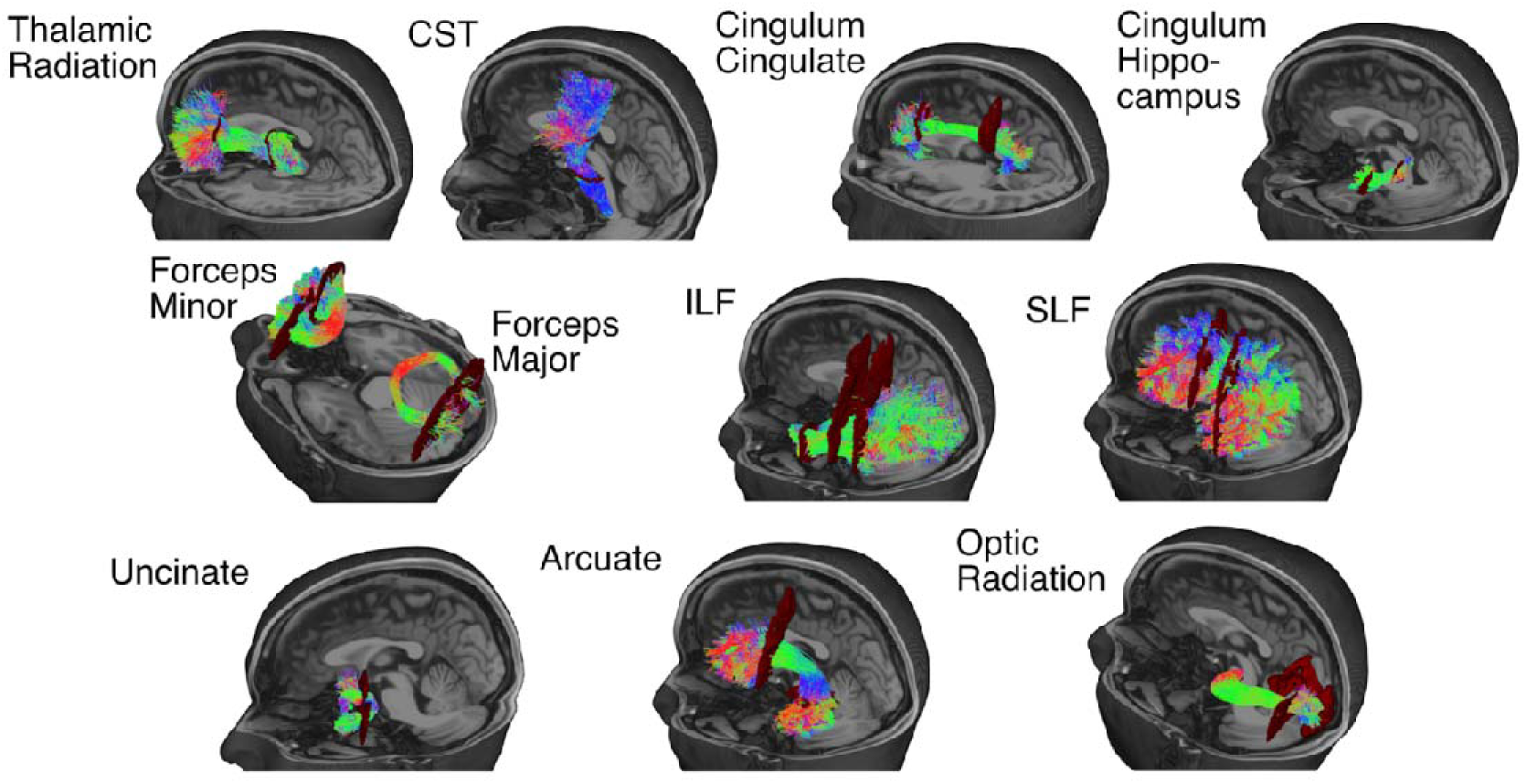
White matter tracts identified from a representative dMRI dataset. White matter tracts were identified from one dMRI dataset acquired from a representative subject (subject 1; Verio A, first run). An automated pipeline (RTP2; Lerma-Usabiaga et al., 2023) was used to identify each tract, shown as colored streamlines, based on regions of interest, shown as dark red volumes. Except for the callosal tracts, namely the forceps minor and forceps major, only the left hemisphere is displayed for visual clarity, although each tract was identified bilaterally for analysis. *CST: corticospinal tract*; *ILF: inferior longitudinal fasciculus*; *SLF: superior longitudinal fasciculus*.

*Voxelwise tissue characterization:* White matter tissue properties in each voxel were characterized using six metrics: fractional anisotropy (FA), MD, intracellular volume fraction (ICVF), orientation dispersion index (ODI), isotropic volume fraction (IsoV), and the T1w/b0 ratio. First, a diffusion tensor model (Basser et al., 1994; Basser & Pierpaoli, 1996) was fitted to dMRI data using a weighted linear least squares method implemented in MRtrix3 (Veraart et al., 2013) to obtain FA and MD. Second, the NODDI model (Zhang et al., 2012) was fitted to dMRI data using the NODDI MATLAB toolbox (http://mig.cs.ucl.ac.uk/index.php?n=Tutorial.NODDImatlab) to obtain ICVF, ODI, and IsoV. The resulting maps were integrated into the RTP2 pipeline as additional input files for tractometry. Third, the T1w/b0 ratio was calculated as the ratio of T1w image intensity to the intensity of the b = 0 s/mm^2^ dMRI image and was used as a proxy for quantitative R1 (Moskovich et al., 2024). To calculate this ratio, T1w images acquired in each run were first co-registered to the T1w image acquired on Verio A during the first run. A T1w/b0 ratio map was then generated by dividing the T1w image by the averaged b = 0 s/mm² image from the preprocessed dMRI data resampled into T1w space. The resulting T1w/b0 ratio image was included as an additional input file for tractometry. For the Tamagawa University PrismaFit A dataset, T1w/b0 was derived only from the first run because no second T1w image was acquired. T1w/b0 could not be calculated for the BCBL PrismaFit B dataset because no MP-RAGE image was acquired.

*Tractometry:* Finally, tractometry was performed by calculating along-tract profiles of white matter tissue properties. To generate tract profiles, the central location of all streamlines in each tract was identified and divided into 100 equally spaced nodes. Tissue properties at each node were summarized by calculating a weighted average of the diffusion properties of the streamlines within that node (Yeatman et al., 2012). Each streamline was weighted according to its Mahalanobis distance from the tract core. Along-tract profiles were generated for each identified tract using the Vistasoft software package (https://github.com/vistalab/vistasoft). Tract-wise mean values for each tract were then calculated by averaging values across all 100 nodes to obtain a representative summary metric.

### 2.4. Harmonization

We investigated the impact of ComBat harmonization (Fortin et al., 2017, 2018; Johnson et al., 2007) on tractometry reproducibility within the same scanner, across different scanners, and across scanner models. Briefly, ComBat was developed to correct for batch effects, which are non-biological factors that introduce unwarranted systematic shifts in measurements (Johnson et al., 2007). In neuroimaging, these effects typically appear as scanner-specific biases that can obscure true biological signals.

ComBat was applied to the tract-wise mean tissue properties, including FA, MD, ICVF, ODI, IsoV, and T1w/b0. These values were calculated by averaging the values across 100 nodes for each tract. Harmonization was performed using publicly available MATLAB code (https://github.com/Jfortin1/ComBatHarmonization/tree/master; Fortin et al., 2017, 2018). Specifically, the batch effects (y, δ) were first estimated using empirical Bayes estimates according to the following formula:

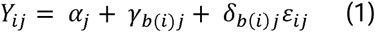

where Y_ij_ is the observed tissue property for subject i and tract J, a_j_ is the averaged value for all subjects of tract J, y_b(i)j_ is the batch effect for batch b(i) and tract J, o_b(i)j_ is the scale factor for batch b(i) and tract J, and ε_ij_ is the random error term representing individual differences, which was assumed to be normally distributed. The batch variable was defined according to the experimental condition, namely scanner and/or protocol. Because the number of subjects in this study was small, covariates were not included in the model to avoid overfitting and unstable parameter estimation (Orlhac et al., 2022).

We then calculated tissue property for subject i and tract J corrected for batch effect Ŷ_ij_ by using the following formula:

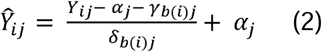

ComBat harmonization was not applied to the PrismaFit B dataset because it included data from only one subject.

### 2.5. Reproducibility and generalizability of tractometry

We evaluated within-scanner reproducibility by comparing tractometry data acquired from the same subject using the same scanner on separate days. Specifically, we compared tract-wise mean values for DTI metrics (FA and MD), NODDI metrics (ICVF and ODI), and T1w/b0 across 20 tracts. Reproducibility and generalizability were quantified using the ICC (2,1), based on a two-way random-effects model with absolute agreement for single measurements (Shrout & Fleiss, 1979). ICC values range from 0, indicating no agreement, to 1, indicating perfect agreement. ICC was calculated separately for each subject across the 20 tracts for each metric, and the resulting values were averaged across the five subjects to obtain a summary measure of within-scanner reproducibility.

To evaluate tractometry generalizability across scanners and scanner models, we calculated ICC using the same procedure. Data from Verio A and Verio B were compared to assess generalizability across scanners of the same model, whereas data from Verio and PrismaFit scanners were compared to assess generalizability across scanner models. For each metric, ICC was calculated for all possible dataset combinations, and the results were visualized in color-coded tables. This analysis was performed using both raw and ComBat-harmonized tractometry data to evaluate the effect of harmonization on reproducibility and generalizability.

The PrismaFit B dataset was available for only one subject, subject 2. Therefore, comparisons involving this dataset are reported separately as single-subject results in the Supplementary Information (Supplementary Figures 3–7).

### 2.6. Minimum detectable change (MDC) analysis for each tract

The preceding analyses evaluated the overall reproducibility and generalizability of tractometry using traveling-head dMRI data. However, the sensitivity of dMRI metrics to hardware and protocol variation is likely to differ by tract. To examine this tract-specific variability, we calculated the minimum detectable change (MDC) for each individual tract using variance estimates derived from pooled datasets across scanners and protocols. This approach allowed us to quantify the vulnerability of specific anatomical structures to cross-scanner and cross-protocol variability and to establish tract-specific reliability thresholds for multisite data. These MDC values provide a benchmark for determining whether observed differences in multisite datasets fall within the expected range of intersite measurement error or may reflect true biological differences.

Specifically, the MDC was calculated at the 95% confidence level (MDC_95_) for each tract. MDC_95_ represents the smallest change that is likely to reflect true biological change rather than measurement error. It was derived from the standard error of measurement (SEM) using the following formula (Haley & Fragala-Pinkham, 2006):

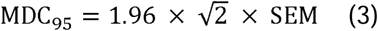

The SEM was derived as:

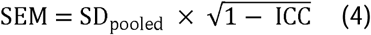

Here, SD_pooled_ represents the pooled standard deviation of the dMRI metric across datasets from all subjects, scanners, and acquisition parameters. Mathematically, it was derived from the square root of the total variance components, including subject, scanner, and residual error, computed during ICC estimation.

Following established practices for quantifying measurement error in dMRI (Schilling et al., 2021), we designed a multisite benchmark to estimate the error threshold for pooled data, such as data from multicenter clinical trials. For this analysis, we used the absolute agreement coefficient, ICC(2,1), across all scanners because it accounts for both random noise and systematic scanner bias (Shrout & Fleiss, 1979). The multisite MDC_95_ was calculated using both raw and ComBat-harmonized tractometry data to quantify the reduction in measurement error achieved by harmonization.

## 3. Results

### 3.1. Within-scanner reproducibility and generalizability across different scanners and scanner models

We collected dMRI datasets from traveling subjects using an experimental design that enabled the evaluation of within-scanner reproducibility and generalizability across scanners and scanner models (Figure 1). We then performed tractometry analysis. Figure 3 shows representative tractometry results for the left corticospinal tract in a single subject. Although the spatial profile of MD along the left corticospinal tract was broadly consistent across datasets, systematic differences in MD values were observed. For quantitative comparisons, tract-wise mean values were calculated for each tract and metric by averaging values along the entire tract trajectory (Figure 3, bottom). These summary measures were then used to evaluate tractometry reproducibility and generalizability across datasets.

**Figure 3.**
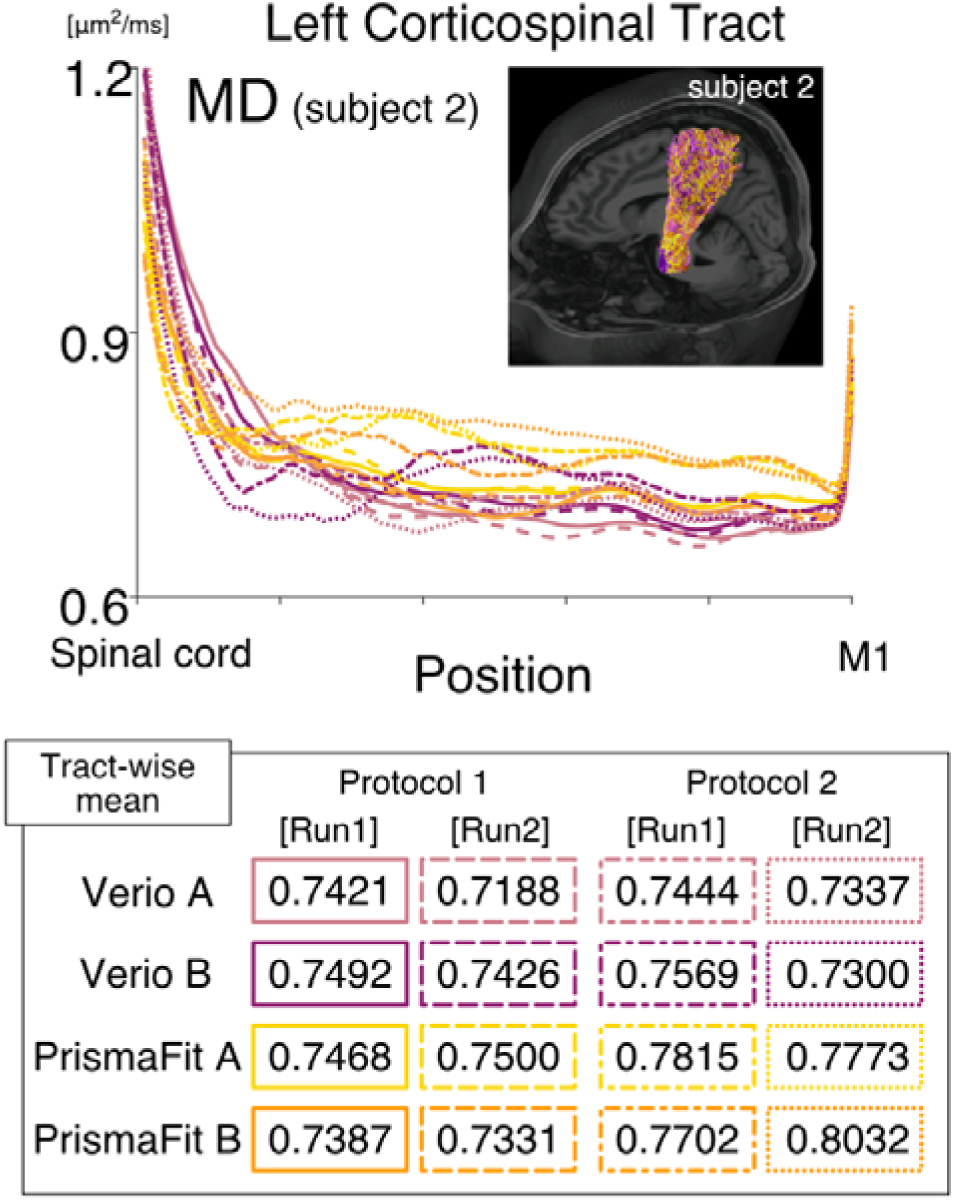
Example tractometry data from the study dataset. *Top panel*: Tract profile of the left corticospinal tract (CST) in a representative subject (subject 2). The upper right image shows the left CST overlaid on axial and sagittal sections of the T1w image. CST streamlines identified from dMRI data acquired using different scanners are superimposed: Verio A, pink; Verio B, purple; PrismaFit A, yellow; PrismaFit B, orange. The plot shows the mean diffusivity (MD) profile along the left CST, with position shown on the horizontal axis and MD on the vertical axis (μm²/ms). Tract profiles derived from each dataset acquired from subject 2 are shown as single curves. Curve styles and colors indicate the dataset and correspond to the box outlines in the bottom panel. *Bottom panel*: Tract-wise mean MD along the left CST in subject 2 for each dataset, obtained by averaging MD across all nodes.

We quantified the reproducibility and generalizability of tract-wise mean values using ICC. MD is presented first as a representative metric. As shown in the left panel of Figure 4, MD showed excellent reproducibility between runs acquired using the same protocol (protocol 1) and the same scanner (Verio A; mean ICC across subjects: 0.990). In contrast, ICC decreased when datasets were acquired from the same subjects using the same protocol but different scanners of the same model (Verio A vs. Verio B; Figure 4, middle panel; mean ICC: 0.949). ICC decreased further when datasets acquired using different scanner models were compared (Verio and PrismaFit; Figure 4, right panel; mean ICC: 0.831). These results indicate that tractometry generalizability declined as instrumental conditions became increasingly different, with the largest differences observed across scanner models.

**Figure 4.**
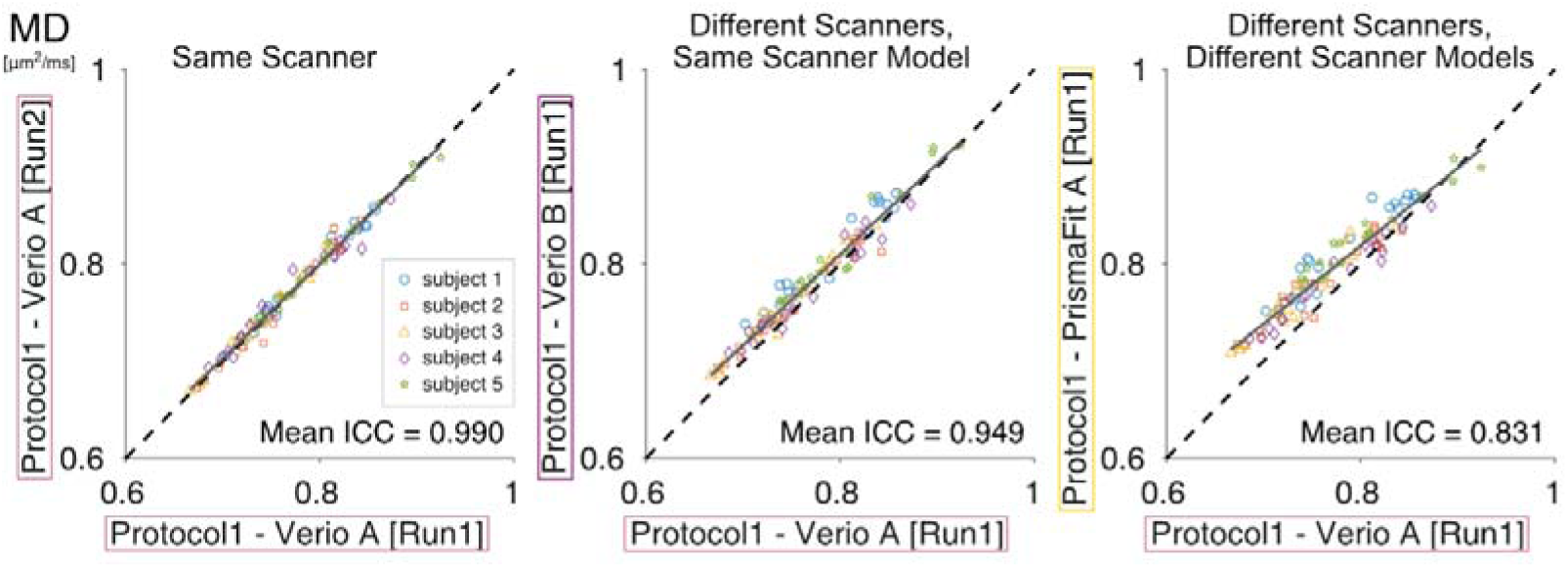
Comparison of MD within a scanner, across scanners of the same model, and across different scanner models using the same acquisition protocol. All comparisons were performed using protocol 1. *Left panel*: Comparison of tractometry data between two datasets acquired using the same scanner, Verio A. Each dot represents the tract-wise mean MD for one of the 20 major white matter tracts. Dot color indicates the subject. The dashed line indicates the identity line (y = x), and the gray solid line indicates the regression line. *Middle panel*: Comparison of tractometry data between datasets acquired using two scanners of the same model, Verio A and Verio B. *Right panel*: Comparison of tractometry data between datasets acquired using different scanner models, Verio A and PrismaFit A. The intraclass correlation coefficient (ICC) was calculated for each subject based on data from 20 tracts, and the mean ICC across subjects is displayed in each panel.

Other dMRI-based metrics, including FA, ICVF, ODI, and IsoV, showed a similar pattern to MD: ICC decreased when data from different scanners were compared and declined more markedly when data from different scanner models were compared (Supplementary Figure 1A–D). Notably, FA, ICVF, and ODI showed higher overall reproducibility and generalizability than MD.

Finally, we examined the reproducibility of T1w/b0, a recently proposed semi-quantitative metric (Moskovich et al., 2024). T1w/b0 showed high ICC when data were acquired using the same scanner (Supplementary Figure 1E; mean ICC: 0.973). ICC remained relatively high across scanners of the same model (mean ICC: 0.950) but decreased markedly when data from different scanner models were compared (mean ICC: 0.271). These results suggest that T1w/b0 is a reliable semi-quantitative metric when data are acquired under similar instrumental conditions and protocols, but its generalizability across scanner models is limited, likely because it depends on both dMRI and MP-RAGE acquisitions.

### 3.2. Generalizability of tractometry across two acquisition protocols

We next examined the effect of acquisition protocol on tractometry generalizability by comparing datasets acquired from the same scanner using two protocols with different acquisition parameters (protocols 1 and 2; Table 1). Figure 5 shows tractometry data for MD from datasets acquired using Verio A as a representative result. As shown in the left and middle panels of Figure 5, MD showed high ICC for both protocols, although the value for protocol 1 was higher than that for protocol 2 (mean ICC: 0.990 and 0.930 for protocols 1 and 2, respectively). However, even when the same scanner was used, ICC decreased when datasets acquired using different protocols were compared (Figure 5, right panel; mean ICC: 0.707). This overall pattern was also observed for other metrics (Supplementary Figure 2). These results indicate that tractometry is sensitive not only to hardware differences but also to differences in acquisition protocols.

**Figure 5.**
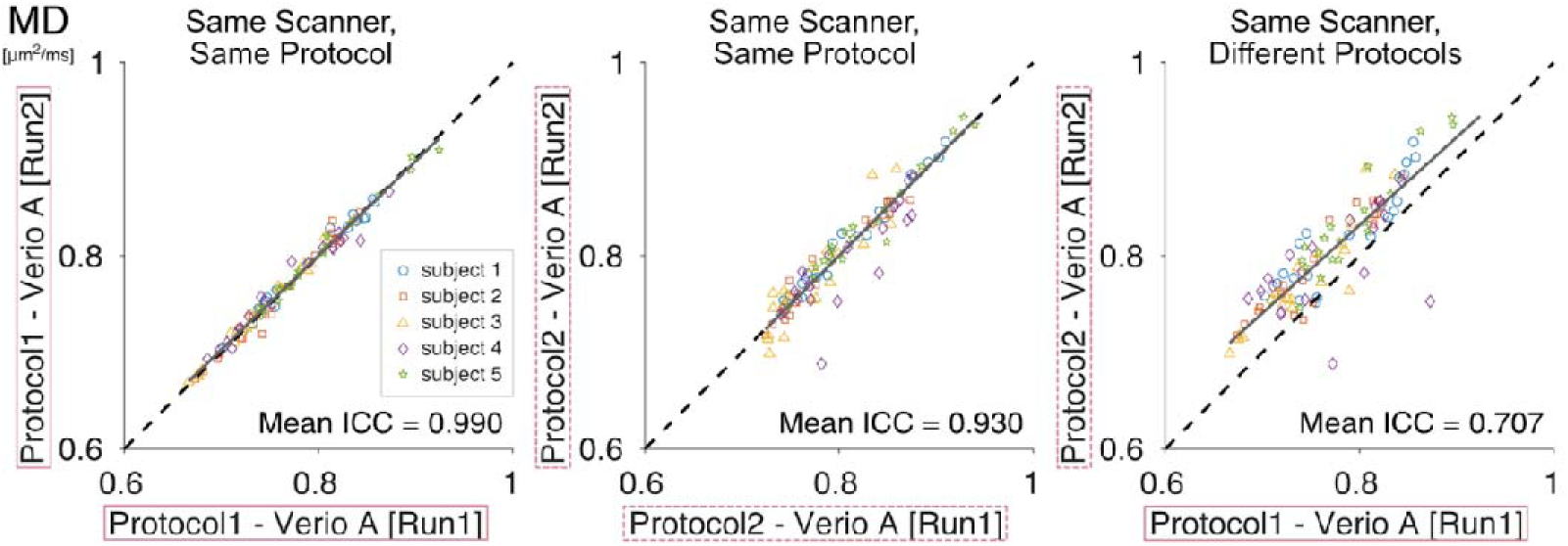
Comparison of MD across datasets acquired using the same scanner with the same or different acquisition protocols. *Left and middle panels*: Comparison of datasets acquired using the same scanner, Verio A, and the same protocol: protocol 1 in the *left panel* and protocol 2 in the *middle panel*. *Right panel*: Comparison of datasets acquired using the same scanner, Verio A, but different protocols, with protocol 1 shown on the horizontal axis and protocol 2 shown on the vertical axis. All other conventions are identical to those in Figure 4.

### 3.3. Impact of ComBat harmonization

To mitigate systematic biases introduced by instrumental and protocol differences, previous studies have proposed ComBat, a harmonization method originally developed for microarray analysis and later adapted for neuroimaging (Fortin et al., 2017, 2018). Here, we evaluated the effect of ComBat harmonization on tractometry reproducibility and generalizability in a dataset in which acquisition conditions, including scanners and protocols, differed incrementally across comparisons.

Figure 6A shows the effects of instrumental and protocol differences on MD tractometry. Before harmonization, ICC was high for datasets acquired using the same scanner (Figure 6A, top left panel; mean ICC: 0.990), lower for datasets acquired using different scanner models with the same protocol (top middle panel; mean ICC: 0.831), and lowest for datasets acquired using different scanner models and different protocols (top right panel; mean ICC: 0.481). When both scanner model and protocol differed, MD measurements showed systematic shifts, causing data points to deviate from the identity line.

**Figure 6.**
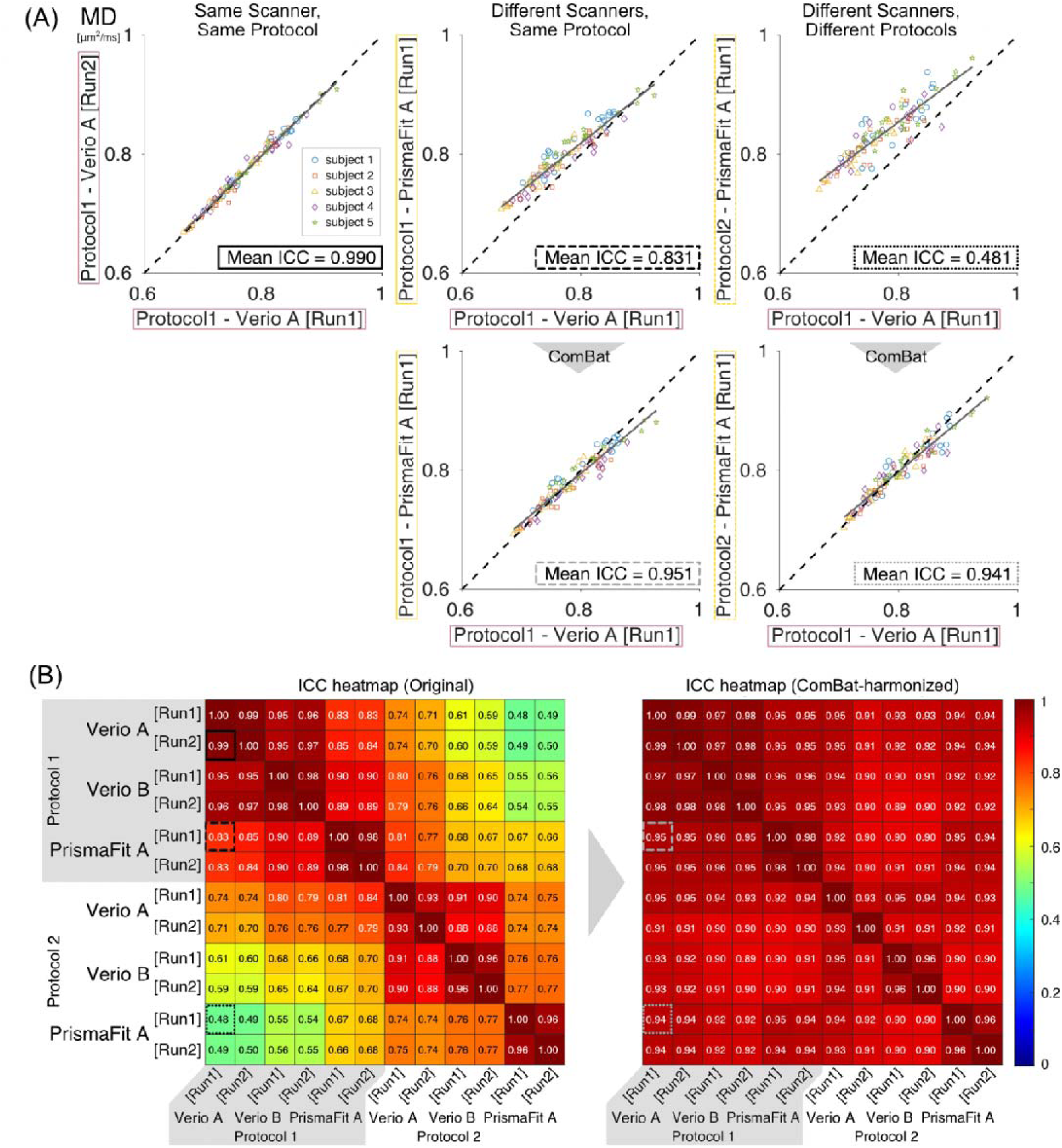
Intraclass correlation coefficient (ICC) of MD across all dataset combinations before and after ComBat harmonization. (**A**) *Top row*: Scatter plots showing MD along 20 major white matter tracts in five subjects and the mean ICC across subjects. *Bottom row*: Corresponding plots after ComBat harmonization. Axes and subject colors are identical to those in the top row. (**B**) Heatmaps showing mean ICC values for MD across all pairwise comparisons before ComBat harmonization (left panel) and after ComBat harmonization (right panel). Rectangles highlighted with black (left panel) or gray (right panel) solid and dashed lines correspond to the comparisons shown in panel A.

After ComBat harmonization, these systematic differences were reduced, and data points were more closely aligned with the identity line (Figure 6A, bottom panel). ICC values improved substantially for comparisons across different scanner models using the same protocol (bottom middle panel; mean ICC: 0.951) and for comparisons across different scanner models and protocols (bottom right panel; mean ICC: 0.941). However, these values did not reach the level observed for datasets acquired using the same scanner, which already showed high ICC before ComBat harmonization (top left panel; mean ICC: 0.990). These results indicate that ComBat harmonization improved tractometry generalizability across scanners and protocols.

Figure 6B (left panel) shows the mean ICC for MD across all dataset combinations before ComBat harmonization (see Supplementary Figures 3–7 for single-subject results, including PrismaFit B, across multiple metrics). Across all scanners, mean ICC values were high when the same scanner and protocol were used (range: 0.93–0.99). Mean ICC values progressively decreased as instrumental and protocol differences increased. Notably, protocol differences had a larger effect than instrumental differences in this dataset. For example, when datasets acquired using protocol 1 were compared, mean ICC ranged from 0.83 to 0.90 despite differences in scanner model (Verio vs. PrismaFit), suggesting that tractometry generalizability remained relatively high despite instrumental differences. In contrast, comparisons of datasets acquired using different protocols on the same scanner showed lower mean ICC values, ranging from 0.64 to 0.74. These results suggest that, in this dataset, protocol consistency was more important for tractometry generalizability than instrumental consistency.

Figure 6B (right panel) shows the mean ICC values after ComBat harmonization. ComBat improved tractometry generalizability across datasets, with all comparisons yielding mean ICC values of 0.89 or higher. These results indicate that systematic variation in tractometry results arising from instrumental and protocol differences can be mitigated using ComBat.

We also evaluated mean ICC values for other tissue-property metrics with and without ComBat harmonization. FA showed relatively higher mean ICC values than MD, even in comparisons across datasets acquired using different protocols (mean ICC ≥ 0.86 in all comparisons; Supplementary Figure 8). Nevertheless, mean ICC increased further after ComBat harmonization (mean ICC ≥ 0.97 in all comparisons; Supplementary Figure 8). When datasets were acquired using protocol 1, FA already showed very high ICC before ComBat harmonization (mean ICC ≥ 0.98 in all comparisons), leaving little room for improvement. ICVF showed a pattern similar to that of MD: mean ICC was affected by both scanner and protocol differences but improved after ComBat harmonization (mean ICC ≥ 0.94 in all comparisons; Supplementary Figure 9). ODI showed a pattern similar to that of FA, with high mean ICC values both before ComBat harmonization (mean ICC ≥ 0.97 when protocol 1 was used) and after ComBat harmonization (mean ICC ≥ 0.97 in all comparisons; Supplementary Figure 10). IsoV showed clear protocol dependence. When protocol 1 was used, IsoV showed relatively high mean ICC values (≥ 0.86); however, when protocol 2 was used, mean ICC values ranged from 0.35 to 0.84, suggesting vulnerability to scanner differences (Supplementary Figure 11), as reported previously (Andica et al., 2020; Bouyagoub et al., 2021; Chung et al., 2016; Mueller et al., 2023). Unlike the other metrics, these errors in protocol 2 datasets were not sufficiently mitigated by ComBat (ICC range: 0.61–0.84). For T1w/b0, scanner differences had a larger effect than protocol differences. Mean ICC values between different scanner models (Verio vs. PrismaFit) were very low (0.14–0.37; Supplementary Figure 12). However, these differences were mitigated by ComBat, resulting in high ICC values (≥ 0.91).

### 3.4. MDC across scanners and protocols

In the preceding sections, we evaluated tractometry reproducibility and generalizability by calculating ICC for dataset pairs across all tracts. However, individual tracts may differ in their vulnerability to scanner and protocol effects. To quantify the extent to which tractometry can detect differences between individuals when data are pooled across scanners and protocols, we calculated MDC at the 95% confidence level (MDC_95_) for each tract and metric after pooling data across scanners and acquisition protocols (Figure 7). This analysis was performed using both original tractometry results and ComBat-harmonized results.

**Figure 7.**
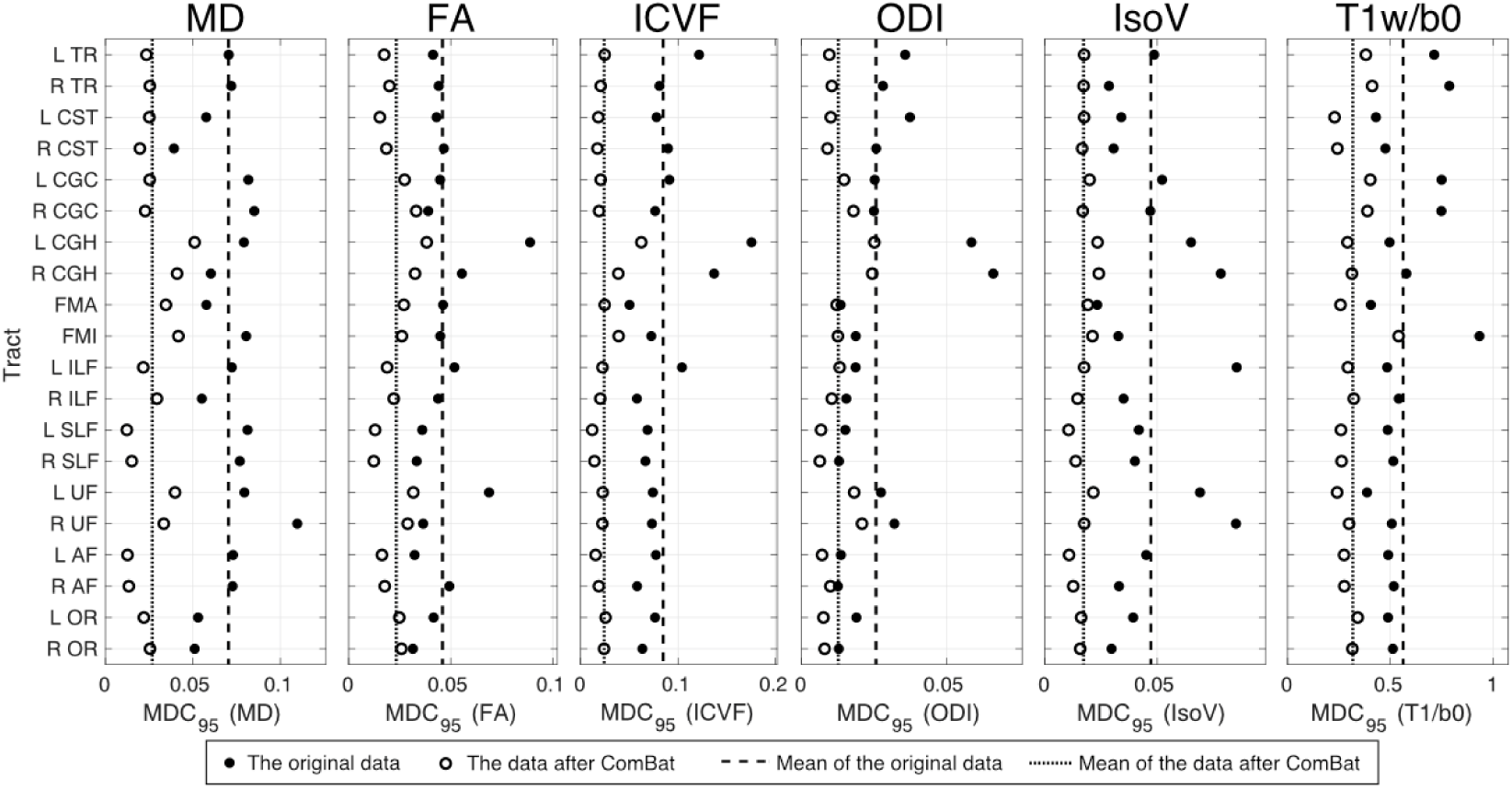
Tract-specific minimum detectable change (MDC_95_) in tractometry data pooled across scanners and protocols. Filled circles indicate MDC_95_ values calculated from the original data, and open circles indicate MDC_95_ values after ComBat harmonization for each tract. L and R indicate the left and right hemispheres, respectively. The dashed line indicates the mean MDC_95_ of the original data across tracts, and the dotted line indicates the mean MDC_95_ after ComBat harmonization. *TR: thalamic radiation, CST: corticosipinal tract, CGC: cingulum cingulate, CGH: cingulum hippocampus, FMA: callosum forceps major, FMI: callosum forceps minor, ILF: inferior longitudinal fasciculus, SLF: superior longitudinal fasciculus, UF: uncinate fasciculus, AF: arcuate fasciculus, OR: optic radiation*.

For MD, MDC_95_ values were broadly similar across most tracts in the original data (mean = 0.070, standard deviation = 0.016). One exception was the right uncinate fasciculus, which showed the largest value (MDC_95_ = 0.11), indicating that MD differences would need to exceed 0.11 to be reliably detected when tractometry data are pooled across scanners and protocols. In contrast, the corticospinal tract and optic radiation showed lower values in both hemispheres (corticospinal tract: MDC_95_ = 0.058 [left], 0.039 [right]; optic radiation: MDC_95_ = 0.053 [left], 0.051 [right]), indicating that tractometry in these tracts was less vulnerable to scanner and protocol differences.

Other metrics showed a pattern broadly similar to that of MD. Some tracts consistently showed larger MDC_95_ values across metrics. In particular, the cingulum hippocampus showed large MDC_95_ values for FA, ICVF, ODI, and IsoV in both hemispheres. The uncinate fasciculus also showed relatively large MDC_95_ values for MD, FA, and IsoV, whereas the left ILF showed a larger MDC_95_ value for IsoV. For T1w/b0, larger MDC_95_ values were observed in the thalamic radiation, forceps minor, and cingulum cingulate. These results suggest that some tracts are generally more vulnerable to scanner and protocol differences, although vulnerability patterns differed by metric.

After ComBat harmonization, MDC_95_ values were consistently reduced, indicating that harmonized tractometry data had a greater ability to detect smaller differences. However, the magnitude of this reduction varied across tracts. Even after ComBat harmonization, the cingulum hippocampus, and uncinate fasciculus retained relatively large MDC_95_ values (Figure 7). This finding suggests that tractometry results in these tracts contained substantial variability that was not fully mitigated by ComBat, likely reflecting tract-specific measurement instability rather than scanner- or protocol-dependent systematic bias alone.

## 4. Discussion

In this study, we quantified the effects of MRI hardware and acquisition protocol differences on tractometry reproducibility and generalizability. Tractometry showed high reproducibility when the same scanner and protocol were used, indicating that it is reliable for basic neuroscience studies in which acquisition conditions can be controlled. However, tractometry became less generalizable as instrumental and protocol differences increased. The magnitude of this systematic bias varied across tissue-property metrics: FA and ODI were relatively robust to scanner and protocol differences, whereas MD and IsoV were more susceptible to these factors. ComBat harmonization mitigated these systematic biases, although post-harmonization ICC values remained below those observed for intra-scanner test–retest reliability when the same protocol was used. Finally, MDC analysis showed that some tracts were more robust than others when data acquired using different scanners and protocols were pooled. Together, these findings provide practical guidance for tractometry studies that require integration of datasets across diverse scanners and protocols.

### 4.1. Different MRI metrics exhibited different levels of tractometry reproducibility and generalizability

Across all evaluated tissue-property metrics, including MD, FA, ICVF, ODI, IsoV, and T1w/b0, tractometry showed high reproducibility when the same scanner and protocol were used. This finding is consistent with previous reports of high test–retest reliability (Kruper et al., 2021; Lerma-Usabiaga et al., 2023). The present study extends previous work by evaluating a broader range of metrics, including conventional DTI metrics, NODDI metrics, and the recently proposed semi-quantitative T1w/b0 metric (Moskovich et al., 2024), whose tractometry reproducibility has not been extensively examined.

When datasets acquired using different scanners and protocols were compared, tractometry generalizability varied substantially across metrics. This finding indicates that some metrics are more resilient than others to systematic bias arising from instrumental and protocol differences.

FA showed strong generalizability despite scanner and protocol differences (Supplementary Figures 1, 2, and 8), whereas MD was more sensitive to these factors (Figures 4–6). This discrepancy may reflect the nature of FA as a normalized index derived from multiple eigenvalues in the diffusion tensor model. If scanner- or protocol-specific effects influence diffusivity measurements uniformly across directions, these scaling effects may be largely canceled out during FA calculation. In contrast, MD is calculated as the arithmetic mean of the three eigenvalues and is therefore more susceptible to absolute scaling differences caused by instrumental and protocol variation. These results are consistent with previous traveling-head studies showing that FA is more robust than MD across scanners (Fox et al., 2012; Kamagata et al., 2015; Saito et al., 2023; Shahim et al., 2017).

Among NODDI-based metrics, ODI showed higher generalizability despite scanner and protocol variation (Supplementary Figures 1, 2, and 10). By contrast, ICVF and, particularly, IsoV showed lower generalizability under heterogeneous acquisition conditions (Supplementary Figures 1, 2, 9, and 11). Because ODI correlates strongly with FA (Zhang et al., 2012), the similar robustness of these two metrics is expected. The robustness of ODI may also reflect its estimation through the Watson distribution, which characterizes relative orientation dispersion rather than absolute diffusivity. This mathematical property may make ODI less sensitive to instrumental and protocol variation, similar to FA. Our findings of high ODI generalizability and greater IsoV sensitivity to acquisition conditions are broadly consistent with previous studies (Bouyagoub et al., 2021; Chung et al., 2016; Lehmann et al., 2021), although some studies have reported different patterns (Andica et al., 2020; Saito et al., 2023), likely because of methodological differences, including ROI selection and acquisition parameters.

Because different TE values were used for Verio and PrismaFit data (Table 1), TE dependence is another plausible contributor to differences in generalizability across dMRI-based metrics. Previous studies have shown that some metrics are affected by TE; for example, prolonged TE can lead to ICVF overestimation, which may partially explain the systematic differences observed between Verio and PrismaFit in this study (Gong et al., 2020; Qin et al., 2009).

Finally, T1w/b0 was highly sensitive to differences in scanner models, particularly Verio versus PrismaFit comparisons (Supplementary Figures 1 and 12). This susceptibility likely arises because T1w/b0 integrates signals from both dMRI and T1w images, each of which is influenced by scanner-specific signal and contrast characteristics. Nevertheless, our results suggest that these scanner model effects were systematic and could be effectively corrected using ComBat harmonization (Supplementary Figure 12B).

### 4.2. Related studies

Numerous traveling-head studies have evaluated dMRI reproducibility; however, analytical approaches have varied considerably. These approaches include ROI-based analysis (Andica et al., 2020; Fox et al., 2012; Grech-Sollars et al., 2015), template- or skeleton-based analysis (Kamagata et al., 2015; Palacios et al., 2017), fixel-based analysis (Mito et al., 2026; Zou et al., 2025), and network analysis (Kurokawa et al., 2021).

Unlike these approaches, tractometry quantifies tissue properties along anatomically defined white matter tracts, supporting anatomically grounded interpretation of dMRI data (Gerig et al., 2004; Jones et al., 2005; Takemura et al., 2024; Yeatman et al., 2012; Yendiki et al., 2011). Because tractometry identifies white matter tracts using prior anatomical information, it can yield robust results without relying on manual ROI selection. Several tractometry software packages are now available (Kruper et al., 2021, 2025; Lerma-Usabiaga et al., 2023; Yeatman et al., 2012; Yendiki et al., 2011), and tractometry has become an established approach for dMRI analysis. Previous studies have demonstrated high test–retest reproducibility when the same scanner and protocol were used (Kamagata et al., 2015; Kruper et al., 2021; Lerma-Usabiaga et al., 2023). The key contribution of the present study is the use of a traveling-head design to evaluate tractometry generalizability across scanners and acquisition protocols.

To our knowledge, only a limited number of studies have investigated tractometry generalizability across scanners and acquisition protocols. Kamagata et al. (2015) evaluated dMRI reproducibility across two scanners from the same vendor using multiple analytical approaches, including tractometry. Using conventional DTI metrics, they reported similarly high within- and between-scanner reproducibility, quantified using coefficients of variation. By including metrics beyond DTI and evaluating two multi-shell high-angular-resolution protocols, our study extends the observations of Kamagata et al. (2015), which were based on relatively low-angular-resolution data with 32 diffusion directions. Schilling et al. (2021) evaluated the extent to which tractometry using FA and MD in seven bilateral white matter tracts depends on scanner, acquisition parameters, and analysis workflow. Similarly, Sierhej et al. (2026) examined the generalizability of DTI and NODDI metrics in four bilateral white matter tracts across three scanners of the same model using the same acquisition protocol. Our study extends these works by quantifying tractometry reproducibility and generalizability for FA, MD, NODDI metrics, and T1w/b0; evaluating all combinations of progressively different acquisition conditions, including the same scanner, different scanners of the same model, different scanner models, and different acquisition protocols; and covering a broader range of tracts.

Chandio et al. (2024) examined tractometry in dMRI datasets acquired using seven different protocols and processed with the BUAN pipeline (Chandio et al., 2024). They analyzed 38 white matter tracts using DTI metrics and proposed node-wise ComBat harmonization along each tract. This approach improved disease detectability in Alzheimer’s disease datasets compared with unharmonized data. Although that study demonstrated the benefit of harmonization for tractometry sensitivity, its primary focus was enhancing group-level pathological detectability rather than evaluating individual-level generalizability because it did not use a traveling-head design. Therefore, our findings complement the observations of Chandio et al. (2024) by directly quantifying tractometry generalizability and the impact of ComBat harmonization at the individual-subject level.

### 4.3. Considerations regarding protocol differences

In addition to showing the sensitivity of tractometry to protocol differences (Figure 5), our results demonstrated that protocol 1 had higher generalizability than protocol 2 across all dMRI metrics (Figure 6; Supplementary Figures 8–12). This difference may be explained by several acquisition-parameter differences between the two protocols. First, protocol 1 included a larger number of diffusion directions, which may improve the robustness of voxelwise model fitting and provide a better signal-to-noise ratio (SNR) for dMRI metrics (Zhan et al., 2010). Second, protocol 1 used a larger voxel size, which may further improve SNR. Third, although protocol 2 is advantageous for reducing susceptibility-induced distortions by shortening TE, its use of parallel imaging and partial Fourier acquisition may reduce SNR (Hamilton et al., 2017), resulting in lower generalizability.

However, establishing universal acquisition-protocol standards remains challenging because each protocol involves tradeoffs. For example, dMRI data acquired using protocol 1 may contain residual distortions even after susceptibility-induced distortion correction. Such residual distortions may adversely affect analyses that require accurate co-registration with structural images. By contrast, protocol 2 has a shorter acquisition time, making it more advantageous for studies involving specific clinical populations or children.

### 4.4. MDC of tractometry across scanners and protocols varies across tracts

Our results also showed that different tracts had different MDC_95_ values when dMRI data acquired using multiple scanners and protocols were pooled (Figure 7). Although ComBat harmonization generally reduced MDC_95_, tract-level differences persisted. These findings indicate that detection sensitivity differs across tracts when tractometry data are pooled across scanners and protocols. These differences may reflect both tract-specific vulnerability to scanner and protocol effects and the inherent instability of dMRI measurements along specific tracts.

Several factors should be considered when interpreting these tract-level differences. The first is tract volume. Smaller tracts are generally more difficult to identify reliably and are more susceptible to partial volume effects from neighboring tracts and tissues. For example, the cingulum hippocampus showed higher MDC_95_, likely because it is a smaller portion of the cingulum bundle and is substantially more difficult to identify. Because data acquired from different scanners and protocols may have different effective spatial resolutions, scanner and protocol effects may have a larger impact on tractometry measurements of this tract. The second factor is susceptibility-induced distortion. In general, dMRI data near the paranasal sinuses or petrous apex/mastoid air complex often exhibit greater susceptibility-induced distortion. Although distortion-correction algorithms can compensate for these effects, residual distortions or signal dropouts may affect dMRI metrics along specific tracts (Takemura et al., 2023). Therefore, because scanners have different gradient strengths and minimum TE values, scanner effects may be larger for tracts near these regions. For example, the uncinate fasciculus is located relatively close to these areas, and its higher variability may partly reflect this factor.

### 4.5. Future directions

We confirmed that tractometry reproducibility for major white matter tracts was very high when the same scanner and protocol were used, consistent with previous findings (Kruper et al., 2021). This result supports the use of tractometry as a quantitative tool for neuroscience studies in which acquisition conditions can be controlled. At the same time, our findings demonstrate that tractometry depends substantially on scanner and protocol choices. Quantifying tractometry generalizability is therefore an essential step toward the future use of tractometry in clinical studies, where dMRI data are often collected in heterogeneous acquisition environments.

Although this study provides a comprehensive evaluation of tractometry reproducibility and generalizability, it could not compare all possible instrumental and acquisition conditions, as no single study can test all combinations. For example, this study could not determine how vendor differences affect tractometry. However, recent developments in vendor-neutral pulse sequence design (Karakuzu et al., 2022) and integrated tractometry software ecosystems (Kruper et al., 2025) may make such comparisons more feasible. Continued efforts to evaluate reproducibility, together with open science practices, will help establish tractometry as a reliable quantitative tool for both neuroscience and clinical studies.

## 5. Conclusion

This study demonstrates that tractometry is highly reproducible when the same scanner and acquisition protocol are used, but its generalizability decreases when scanners, scanner models, or acquisition protocols differ. These findings highlight the importance of quantifying and accounting for scanner- and protocol-dependent effects, particularly in multisite studies that integrate heterogeneous dMRI datasets. Furthermore, the MDC of tractometry data pooled across scanners and protocols varied across metrics and tracts, indicating that sensitivity to scanner and protocol effects is metric- and tract-dependent.

## Data and code availability

We plan to make code for replicating figures and statistical analysis in GitHub, upon the acceptance of the manuscript ([URL will be inserted upon acceptance]). In addition, we also plan to release the preprocessed and anonymized dataset after skull-stripping publicly available in a public repository ([URL will be inserted upon acceptance]).

## Acknowledgment

We thank Editage (http://www.editage.com) for editing and reviewing this manuscript for the English language.

## Author contributions

Conceptualization: DT T. Matsuda GLU HT, Data curation: DT IY TK ST T. Miyata GLU HT, Formal analysis: DT GLU, Funding acquisition: DT T. Matsuda GLU HT, Investigation: DT, Project administration: T. Matsuda GLU HT, Resources: IY T. Matsuda GLU HT, Software: GLU, Supervision: GLU HT, Validation: DT GLU HT, Visualization: DT, Writing—Original draft preparation: DT GLU HT, Writing—Review & editing: DT IY TK ST T. Miyata T. Matsuda GLU HT.

## Disclosure of the use of artificial intelligence

The authors utilized Gemini 3 Flash and Grammarly for linguistic editing and grammatical refinement. Following the AI-assisted processing, the manuscript was manually revised by the authors to ensure accuracy and to reflect the authors’ original perspectives. The final content is the sole responsibility of the authors.

## References

Alagapan, S., Choi, K. S., Heisig, S., Riva-Posse, P., Crowell, A., Tiruvadi, V., Obatusin, M., Veerakumar, A., Waters, A. C., Gross, R. E., Quinn, S., Denison, L., O’Shaughnessy, M., Connor, M., Canal, G., Cha, J., Hershenberg, R., Nauvel, T., Isbaine, F., … Rozell, C. J. (2023). Cingulate dynamics track depression recovery with deep brain stimulation. Nature, 622(7981), 130–138.

Amemiya, K., Naito, E., & Takemura, H. (2021). Age dependency and lateralization in the three branches of the human superior longitudinal fasciculus. Cortex, 139, 116–133.

Andersson, J. L. R., Skare, S., & Ashburner, J. (2003). How to correct susceptibility distortions in spin-echo echo-planar images: application to diffusion tensor imaging. NeuroImage, 20(2), 870–888.

Andersson, J. L. R., & Sotiropoulos, S. N. (2016). An integrated approach to correction for off-resonance effects and subject movement in diffusion MR imaging. NeuroImage, 125, 1063–1078.

Andica, C., Kamagata, K., Hayashi, T., Hagiwara, A., Uchida, W., Saito, Y., Kamiya, K., Fujita, S., Akashi, T., Wada, A., Abe, M., Kusahara, H., Hori, M., & Aoki, S. (2020). Scan-rescan and inter-vendor reproducibility of neurite orientation dispersion and density imaging metrics. Neuroradiology, 62(4), 483–494.

Assaf, Y., Johansen-Berg, H., & Thiebaut de Schotten, M. (2019). The role of diffusion MRI in neuroscience. NMR in Biomedicine, 32(4), e3762.

Bach, M., Laun, F. B., Leemans, A., Tax, C. M. W., Biessels, G. J., Stieltjes, B., & Maier-Hein, K. H. (2014). Methodological considerations on tract-based spatial statistics (TBSS). NeuroImage, 100, 358–369.

Basser, P. J., Mattiello, J., & LeBihan, D. (1994). Estimation of the effective self-diffusion tensor from the NMR spin echo. Journal of Magnetic Resonance. Series B, 103(3), 247–254.

Basser, P. J., & Pierpaoli, C. (1996). Microstructural and physiological features of tissues elucidated by quantitative-diffusion-tensor MRI. Journal of Magnetic Resonance. Series B, 111(3), 209–219.

Benson, N. C., Butt, O. H., Brainard, D. H., & Aguirre, G. K. (2014). Correction of distortion in flattened representations of the cortical surface allows prediction of V1-V3 functional organization from anatomy. PLoS Computational Biology, 10(3), e1003538.

Benson, N. C., Butt, O. H., Datta, R., Radoeva, P. D., Brainard, D. H., & Aguirre, G. K. (2012). The retinotopic organization of striate cortex is well predicted by surface topology. Current Biology, 22(21), 2081–2085.

Boukadi, M., Marcotte, K., Bedetti, C., Houde, J.-C., Desautels, A., Deslauriers-Gauthier, S., Chapleau, M., Boré, A., Descoteaux, M., & Brambati, S. M. (2018). Test-retest reliability of diffusion measures extracted along white matter language fiber bundles using HARDI-based tractography. Frontiers in Neuroscience, 12, 1055.

Bouyagoub, S., Dowell, N. G., Gabel, M., & Cercignani, M. (2021). Comparing multiband and singleband EPI in NODDI at 3 T: what are the implications for reproducibility and study sample sizes? Magma, 34(4), 499–511.

Briggs, F., Kiley, C. W., Callaway, E. M., & Usrey, W. M. (2016). Morphological substrates for parallel streams of corticogeniculate feedback originating in both V1 and V2 of the macaque monkey. Neuron, 90(2), 388–399.

Catani, M., & Ffytche, D. H. (2005). The rises and falls of disconnection syndromes. Brain: A Journal of Neurology, 128(Pt 10), 2224–2239.

Catani, M., Howard, R. J., Pajevic, S., & Jones, D. K. (2002). Virtual in vivo interactive dissection of white matter fasciculi in the human brain. NeuroImage, 17(1), 77–94.

Chandio, B. Q., Risacher, S. L., Pestilli, F., Bullock, D., Yeh, F.-C., Koudoro, S., Rokem, A., Harezlak, J., & Garyfallidis, E. (2020). Bundle analytics, a computational framework for investigating the shapes and profiles of brain pathways across populations. Scientific Reports, 10(1), 17149.

Chandio, B. Q., Villalon-Reina, J. E., Nir, T. M., Thomopoulos, S. I., Feng, Y., Benavidez, S., Jahanshad, N., Harezlak, J., Garyfallidis, E., & Thompson, P. M. (2024). Bundle ANalytics based data harmonization for multi-site diffusion MRI tractometry. Annual International Conference of the IEEE Engineering in Medicine and Biology Society, 2024, 1–7.

Chung, A. W., Seunarine, K. K., & Clark, C. A. (2016). NODDI reproducibility and variability with magnetic field strength: A comparison between 1.5 T and 3 T. Human Brain Mapping, 37(12), 4550–4565.

Cousineau, M., Jodoin, P.-M., Morency, F. C., Rozanski, V., Grand’Maison, M., Bedell, B. J., & Descoteaux, M. (2017). A test-retest study on Parkinson’s PPMI dataset yields statistically significant white matter fascicles. NeuroImage: Clinical, 16, 222–233.

Fischl, B. (2012). FreeSurfer. NeuroImage, 62(2), 774–781.

Forkel, S. J., Friedrich, P., Thiebaut de Schotten, M., & Howells, H. (2022). White matter variability, cognition, and disorders: a systematic review. Brain Structure and Function, 227(2), 529–544.

Fortin, J.-P., Cullen, N., Sheline, Y. I., Taylor, W. D., Aselcioglu, I., Cook, P. A., Adams, P., Cooper, C., Fava, M., McGrath, P. J., McInnis, M., Phillips, M. L., Trivedi, M. H., Weissman, M. M., & Shinohara, R. T. (2018). Harmonization of cortical thickness measurements across scanners and sites. NeuroImage, 167, 104–120.

Fortin, J.-P., Parker, D., Tunç, B., Watanabe, T., Elliott, M. A., Ruparel, K., Roalf, D. R., Satterthwaite, T. D., Gur, R. C., Gur, R. E., Schultz, R. T., Verma, R., & Shinohara, R. T. (2017). Harmonization of multi-site diffusion tensor imaging data. NeuroImage, 161, 149–170.

Fox, R. J., Sakaie, K., Lee, J.-C., Debbins, J. P., Liu, Y., Arnold, D. L., Melhem, E. R., Smith, C. H., Philips, M. D., Lowe, M., & Fisher, E. (2012). A validation study of multicenter diffusion tensor imaging: reliability of fractional anisotropy and diffusivity values. American Journal of Neuroradiology, 33(4), 695–700.

Gerig, G., Gouttard, S., & Corouge, I. (2004). Analysis of brain white matter via fiber tract modeling. Annual International Conference of the IEEE Engineering in Medicine and Biology Society, 2004, 4421–4424.

Glozman, T., Bruckert, L., Pestilli, F., Yecies, D. W., Guibas, L. J., & Yeom, K. W. (2018). Framework for shape analysis of white matter fiber bundles. NeuroImage, 167, 466–477.

Gong, T., Tong, Q., He, H., Sun, Y., Zhong, J., & Zhang, H. (2020). MTE-NODDI: Multi-TE NODDI for disentangling non-T2-weighted signal fractions from compartment-specific T2 relaxation times. NeuroImage, 217(116906), 116906.

Gorgolewski, K. J., Auer, T., Calhoun, V. D., Craddock, R. C., Das, S., Duff, E. P., Flandin, G., Ghosh, S. S., Glatard, T., Halchenko, Y. O., Handwerker, D. A., Hanke, M., Keator, D., Li, X., Michael, Z., Maumet, C., Nichols, B. N., Nichols, T. E., Pellman, J., … Poldrack, R. A. (2016). The brain imaging data structure, a format for organizing and describing outputs of neuroimaging experiments. Scientific Data, 3(1), 160044.

Grech-Sollars, M., Hales, P. W., Miyazaki, K., Raschke, F., Rodriguez, D., Wilson, M., Gill, S. K., Banks, T., Saunders, D. E., Clayden, J. D., Gwilliam, M. N., Barrick, T. R., Morgan, P. S., Davies, N. P., Rossiter, J., Auer, D. P., Grundy, R., Leach, M. O., Howe, F. A., … Clark, C. A. (2015). Multi-centre reproducibility of diffusion MRI parameters for clinical sequences in the brain: Multi-centre reproducibility of diffusion MRI using clinical sequences. NMR in Biomedicine, 28(4), 468–485.

Halchenko, Y. O., Goncalves, M., Ghosh, S., Velasco, P., di Oleggio Castello, M. V., Salo, T., Wodder, J. T., II, Hanke, M., Sadil, P., Gorgolewski, K. J., Ioanas, H.-I., Rorden, C., Hendrickson, T. J., Dayan, M., Houlihan, S. D., Kent, J., Strauss, T., Lee, J., To, I., … Kennedy, D. N. (2025). HeuDiConv — flexible DICOM conversion into structured directory layouts. Zenodo. 10.5281/ZENODO.15080551

Haley, S. M., & Fragala-Pinkham, M. A. (2006). Interpreting change scores of tests and measures used in physical therapy. Physical Therapy, 86(5), 735–743.

Hamilton, J., Franson, D., & Seiberlich, N. (2017). Recent advances in parallel imaging for MRI. Progress in Nuclear Magnetic Resonance Spectroscopy, 101, 71–95.

Iglesias, J. E., Insausti, R., Lerma-Usabiaga, G., Bocchetta, M., Van Leemput, K., Greve, D. N., van der Kouwe, A., Alzheimer’s Disease Neuroimaging Initiative, Fischl, B., Caballero-Gaudes, C., & Paz-Alonso, P. M. (2018). A probabilistic atlas of the human thalamic nuclei combining ex vivo MRI and histology. NeuroImage, 183, 314–326.

Jenkinson, M., Beckmann, C. F., Behrens, T. E. J., Woolrich, M. W., & Smith, S. M. (2012). FSL. NeuroImage, 62(2), 782–790.

Jeurissen, B., Tournier, J.-D., Dhollander, T., Connelly, A., & Sijbers, J. (2014). Multi-tissue constrained spherical deconvolution for improved analysis of multi-shell diffusion MRI data. NeuroImage, 103, 411–426.

Johnson, W. E., Li, C., & Rabinovic, A. (2007). Adjusting batch effects in microarray expression data using empirical Bayes methods. Biostatistics, 8(1), 118–127.

Jones, D. K., Travis, A. R., Eden, G., Pierpaoli, C., & Basser, P. J. (2005). PASTA: pointwise assessment of streamline tractography attributes. Magnetic Resonance in Medicine, 53(6), 1462–1467.

Jovicich, J., Czanner, S., Greve, D., Haley, E., van der Kouwe, A., Gollub, R., Kennedy, D., Schmitt, F., Brown, G., Macfall, J., Fischl, B., & Dale, A. (2006). Reliability in multi-site structural MRI studies: effects of gradient non-linearity correction on phantom and human data. NeuroImage, 30(2), 436–443.

Kamagata, K., Shimoji, K., Hori, M., Nishikori, A., Tsuruta, K., Yoshida, M., Kamiya, K., Irie, R., Suzuki, M., Kyogoku, S., Suzuki, Y., Sato, N., & Aoki, S. (2015). Intersite reliability of diffusion tensor imaging on two 3T scanners. Magnetic Resonance in Medical Sciences, 14(3), 227–233.

Karakuzu, A., Biswas, L., Cohen-Adad, J., & Stikov, N. (2022). Vendor-neutral sequences and fully transparent workflows improve inter-vendor reproducibility of quantitative MRI. Magnetic Resonance in Medicine, 88(3), 1212–1228.

Kellner, E., Dhital, B., Kiselev, V. G., & Reisert, M. (2016). Gibbs-ringing artifact removal based on local subvoxel-shifts. Magnetic Resonance in Medicine, 76(5), 1574–1581.

Koike, S., Tanaka, S. C., Okada, T., Aso, T., Yamashita, A., Yamashita, O., Asano, M., Maikusa, N., Morita, K., Okada, N., Fukunaga, M., Uematsu, A., Togo, H., Miyazaki, A., Murata, K., Urushibata, Y., Autio, J., Ose, T., Yoshimoto, J., … Brain/MINDS Beyond Human Brain MRI Group. (2021). Brain/MINDS beyond human brain MRI project: A protocol for multi-level harmonization across brain disorders throughout the lifespan. NeuroImage. Clinical, 30(102600), 102600.

Kruper, J., Richie-Halford, A., Qiao, J., Gilmore, A., Chang, K., Grotheer, M., Roy, E., Caffarra, S., Gomez, T., Chou, S., Cieslak, M., Koudoro, S., Garyfallidis, E., Satterthwaite, T. D., Yeatman, J. D., & Rokem, A. (2025). A software ecosystem for brain tractometry processing, analysis, and insight. PLoS Computational Biology, 21(8), e1013323.

Kruper, J., Yeatman, J. D., Richie-Halford, A., Bloom, D., Grotheer, M., Caffarra, S., Kiar, G., Karipidis, I. I., Roy, E., Chandio, B. Q., Garyfallidis, E., & Rokem, A. (2021). Evaluating the reliability of human brain white matter tractometry. Aperture Neuro, 1(1), 1–25.

Kurokawa, R., Kamiya, K., Koike, S., Nakaya, M., Uematsu, A., Tanaka, S. C., Kamagata, K., Okada, N., Morita, K., Kasai, K., & Abe, O. (2021). Cross-scanner reproducibility and harmonization of a diffusion MRI structural brain network: A traveling subject study of multi-b acquisition. NeuroImage, 245(118675), 118675.

Lebel, C., Treit, S., & Beaulieu, C. (2019). A review of diffusion MRI of typical white matter development from early childhood to young adulthood. NMR in Biomedicine, 32(4), e3778.

Lehmann, N., Aye, N., Kaufmann, J., Heinze, H.-J., Düzel, E., Ziegler, G., & Taubert, M. (2021). Longitudinal reproducibility of Neurite Orientation Dispersion and Density Imaging (NODDI) derived metrics in the white matter. Neuroscience, 457, 165–185.

Lerma-Usabiaga, G., Liu, M., Paz-Alonso, P. M., & Wandell, B. A. (2023). Reproducible Tract Profiles 2 (RTP2) suite, from diffusion MRI acquisition to clinical practice and research. Scientific Reports, 13(1), 6010.

Lerma-Usabiaga, G., Mukherjee, P., Ren, Z., Perry, M. L., & Wandell, B. A. (2019). Replication and generalization in applied neuroimaging. NeuroImage, 202(116048), 116048.

Liu, M., Lerma-Usabiaga, G., Clascá, F., & Paz-Alonso, P. M. (2022). Reproducible protocol to obtain and measure first-order relay human thalamic white-matter tracts. NeuroImage, 262(119558), 119558.

Li, X., Morgan, P. S., Ashburner, J., Smith, J., & Rorden, C. (2016). The first step for neuroimaging data analysis: DICOM to NIfTI conversion. Journal of Neuroscience Methods, 264, 47–56.

Maier-Hein, K. H., Neher, P. F., Houde, J.-C., Côté, M.-A., Garyfallidis, E., Zhong, J., Chamberland, M., Yeh, F.-C., Lin, Y.-C., Ji, Q., Reddick, W. E., Glass, J. O., Chen, D. Q., Feng, Y., Gao, C., Wu, Y., Ma, J., He, R., Li, Q., … Descoteaux, M. (2017). The challenge of mapping the human connectome based on diffusion tractography. Nature Communications, 8(1), 1349.

Miller, K. L., Alfaro-Almagro, F., Bangerter, N. K., Thomas, D. L., Yacoub, E., Xu, J., Bartsch, A. J., Jbabdi, S., Sotiropoulos, S. N., Andersson, J. L. R., Griffanti, L., Douaud, G., Okell, T. W., Weale, P., Dragonu, I., Garratt, S., Hudson, S., Collins, R., Jenkinson, M., … Smith, S. M. (2016). Multimodal population brain imaging in the UK Biobank prospective epidemiological study. Nature Neuroscience, 19(11), 1523–1536.

Mito, R., Genc, S., Halim, J., Yang, J. Y.-M., Tournier, J.-D., Kean, M., Kokkinos, C., McIntyre, R., Di Biase, M. A., Smith, R. E., & Zalesky, A. (2026). TRAMFIX: TRavelling Across Melbourne for FIXel-based analysis (a reproducibility and reliability study). Imaging Neuroscience, 4, IMAG.a.1153.

Mori, S., & van Zijl, P. C. M. (2002). Fiber tracking: principles and strategies - a technical review. NMR in Biomedicine, 15(7-8), 468–480.

Mori, S., & Zhang, J. (2006). Principles of diffusion tensor imaging and its applications to basic neuroscience research. Neuron, 51(5), 527–539.

Moskovich, S., Shtangel, O., & Mezer, A. A. (2024). Approximating R1 and R2: A quantitative approach to clinical weighted MRI. Human Brain Mapping, 45(18), e70102.

Mueller, C., Goodman, A. M., Nenert, R., Allendorfer, J. B., Philip, N. S., Correia, S., Oster, R. A., LaFrance, W. C., Jr, & Szaflarski, J. P. (2023). Repeatability of neurite orientation dispersion and density imaging in patients with traumatic brain injury. Journal of Neuroimaging, 33(5), 802–824.

Orlhac, F., Eertink, J. J., Cottereau, A.-S., Zijlstra, J. M., Thieblemont, C., Meignan, M., Boellaard, R., & Buvat, I. (2022). A guide to ComBat harmonization of imaging biomarkers in multicenter studies. Journal of Nuclear Medicine, 63(2), 172–179.

Palacios, E. M., Martin, A. J., Boss, M. A., Ezekiel, F., Chang, Y. S., Yuh, E. L., Vassar, M. J., Schnyer, D. M., MacDonald, C. L., Crawford, K. L., Irimia, A., Toga, A. W., Mukherjee, P., & TRACK-TBI Investigators. (2017). Toward precision and reproducibility of diffusion tensor imaging: A multicenter diffusion phantom and traveling volunteer study. American Journal of Neuroradiology, 38(3), 537–545.

Qin, W., Yu, C. S., Zhang, F., Du, X. Y., Jiang, H., Yan, Y. X., & Li, K. C. (2009). Effects of echo time on diffusion quantification of brain white matter at 1.5 T and 3.0 T. Magnetic Resonance in Medicine, 61(4), 755–760.

Saito, Y., Kamagata, K., Andica, C., Maikusa, N., Uchida, W., Takabayashi, K., Yoshida, S., Hagiwara, A., Fujita, S., Akashi, T., Wada, A., Irie, R., Shimoji, K., Hori, M., Kamiya, K., Koike, S., Hayashi, T., & Aoki, S. (2023). Traveling subject-informed harmonization increases reliability of brain diffusion tensor and neurite mapping. Aging and Disease, 15(6), 2770–2785.

Schilling, K. G., Petit, L., Rheault, F., Remedios, S., Pierpaoli, C., Anderson, A. W., Landman, B. A., & Descoteaux, M. (2020). Brain connections derived from diffusion MRI tractography can be highly anatomically accurate-if we know where white matter pathways start, where they end, and where they do not go. Brain Structure and Function, 225(8), 2387–2402.

Schilling, K. G., Tax, C. M. W., Rheault, F., Hansen, C., Yang, Q., Yeh, F.-C., Cai, L., Anderson, A. W., & Landman, B. A. (2021). Fiber tractography bundle segmentation depends on scanner effects, vendor effects, acquisition resolution, diffusion sampling scheme, diffusion sensitization, and bundle segmentation workflow. NeuroImage, 242(118451), 118451.

Shahim, P., Holleran, L., Kim, J. H., & Brody, D. L. (2017). Test-retest reliability of high spatial resolution diffusion tensor and diffusion kurtosis imaging. Scientific Reports, 7(1), 11141.

Sherbondy, A. J., Dougherty, R. F., Napel, S., & Wandell, B. A. (2008). Identifying the human optic radiation using diffusion imaging and fiber tractography. Journal of Vision, 8(10), 12.1–11.

Shrout, P. E., & Fleiss, J. L. (1979). Intraclass correlations: uses in assessing rater reliability. Psychological Bulletin, 86(2), 420–428.

Sierhej, A., Correia, M. M., Evans, C. J, Seunarine, K. K., Clayden, J. D., Smith, N. A. S., Hall, M. G., & Clark, C. A. (2026). Multi-centre reproducibility of DTI and NODDI in white matter tracts segmented using TractFinder across three MRI scanners of the same model. Human Brain Mapping, 47(5), e70491.

Smith, R. E., Tournier, J. D., Calamante, F. & Connelly, A. (2013). SIFT: Spherical-deconvolution informed filtering of tractograms. Neuroimage, 67, 298–312.

Sotiropoulos, S. N., & Zalesky, A. (2019). Building connectomes using diffusion MRI: why, how and but. NMR in Biomedicine, 32(4), e3752.

Takemura, H., Caiafa, C. F., Wandell, B. A., & Pestilli, F. (2016). Ensemble tractography. PLoS Computational Biology, 12(2), e1004692.

Takemura, H., Kruper, J. A., Miyata, T., & Rokem, A. (2024). Tractometry of human visual white matter pathways in health and disease. Magnetic Resonance in Medical Sciences, 23(3), 316–340.

Takemura, H., Liu, W., Kuribayashi, H., Miyata, T., & Kida, I. (2023). Evaluation of simultaneous multi-slice readout-segmented diffusion-weighted MRI acquisition in human optic nerve measurements. Magnetic Resonance Imaging, 102, 103–114.

Thomason, M. E., & Thompson, P. M. (2011). Diffusion imaging, white matter, and psychopathology. Annual Review of Clinical Psychology, 7(1), 63–85.

Tong, Q., He, H., Gong, T., Li, C., Liang, P., Qian, T., Sun, Y., Ding, Q., Li, K., & Zhong, J. (2019). Reproducibility of multi-shell diffusion tractography on traveling subjects: A multicenter study prospective. Magnetic Resonance Imaging, 59, 1–9.

Tong, Q., He, H., Gong, T., Li, C., Liang, P., Qian, T., Sun, Y., Ding, Q., Li, K., & Zhong, J. (2020). Multicenter dataset of multi-shell diffusion MRI in healthy traveling adults with identical settings. Scientific Data, 7(1), 157.

Tournier, J. D., Calamante, F., & Connelly, A. (2010). Improved probabilistic streamlines tractography by 2nd order integration over fibre orientation distributions. Proceedings of the International Society for Magnetic Resonance in Medicine, 18, 1670.

Tournier, J.-D., Smith, R., Raffelt, D., Tabbara, R., Dhollander, T., Pietsch, M., Christiaens, D., Jeurissen, B., Yeh, C.-H., & Connelly, A. (2019). MRtrix3: A fast, flexible and open software framework for medical image processing and visualisation. NeuroImage, 202(116137), 116137.

Veraart, J., Sijbers, J., Sunaert, S., Leemans, A., & Jeurissen, B. (2013). Weighted linear least squares estimation of diffusion MRI parameters: Strengths, limitations, and pitfalls. NeuroImage, 81, 335–346.

Vollmar, C., O’Muircheartaigh, J., Barker, G. J., Symms, M. R., Thompson, P., Kumari, V., Duncan, J. S., Richardson, M. P., & Koepp, M. J. (2010). Identical, but not the same: intra-site and inter-site reproducibility of fractional anisotropy measures on two 3.0T scanners. NeuroImage, 51(4), 1384–1394.

Wakana, S., Caprihan, A., Panzenboeck, M. M., Fallon, J. H., Perry, M., Gollub, R. L., Hua, K., Zhang, J., Jiang, H., Dubey, P., Blitz, A., van Zijl, P., & Mori, S. (2007). Reproducibility of quantitative tractography methods applied to cerebral white matter. NeuroImage, 36(3), 630–644.

Wandell, B. A. (2016). Clarifying human white matter. Annual Review of Neuroscience, 39, 103–128.

Yamashita, A., Yahata, N., Itahashi, T., Lisi, G., Yamada, T., Ichikawa, N., Takamura, M., Yoshihara, Y., Kunimatsu, A., Okada, N., Yamagata, H., Matsuo, K., Hashimoto, R., Okada, G., Sakai, Y., Morimoto, J., Narumoto, J., Shimada, Y., Kasai, K., … Imamizu, H. (2019). Harmonization of resting-state functional MRI data across multiple imaging sites via the separation of site differences into sampling bias and measurement bias. PLoS Biology, 17(4), e3000042.

Yeatman, J. D., Dougherty, R. F., Myall, N. J., Wandell, B. A., & Feldman, H. M. (2012). Tract profiles of white matter properties: automating fiber-tract quantification. PloS One, 7(11), e49790.

Yendiki, A., Panneck, P., Srinivasan, P., Stevens, A., Zöllei, L., Augustinack, J., Wang, R., Salat, D., Ehrlich, S., Behrens, T., Jbabdi, S., Gollub, R., & Fischl, B. (2011). Automated probabilistic reconstruction of white-matter pathways in health and disease using an atlas of the underlying anatomy. Frontiers in Neuroinformatics, 5, 23.

Yukie, M., & Iwai, E. (1981). Direct projection from the dorsal lateral geniculate nucleus to the prestriate cortex in macaque monkeys. The Journal of Comparative Neurology, 201(1), 81–97.

Zhang, H., Schneider, T., Wheeler-Kingshott, C. A., & Alexander, D. C. (2012). NODDI: practical in vivo neurite orientation dispersion and density imaging of the human brain. NeuroImage, 61(4), 1000–1016.

Zhang, W., Olivi, A., Hertig, S. J., van Zijl, P., & Mori, S. (2008). Automated fiber tracking of human brain white matter using diffusion tensor imaging. NeuroImage, 42(2), 771–777.

Zhan, L., Leow, A. D., Jahanshad, N., Chiang, M.-C., Barysheva, M., Lee, A. D., Toga, A. W., McMahon, K. L., de Zubicaray, G. I., Wright, M. J., & Thompson, P. M. (2010). How does angular resolution affect diffusion imaging measures? NeuroImage, 49(2), 1357–1371.

Zou, R., Kamagata, K., Mito, R., Takabayashi, K., Andica, C., Uchida, W., Guo, S., Kitagawa, T., Fujita, S., Uematsu, A., Maikusa, N., Koike, S., Aoki, S., & Alzheimer’s Disease Neuroimaging Initiative. (2025). Utility of harmonisation for fixel-based metrics in travelling subjects and Alzheimer’s disease data. Human Brain Mapping, 46(16), e70408.

