## Supplementary Information for "Tractometry reproducibility and generalizability across scanners, scanner models, and acquisition protocols"

^†^: equal senior contribution

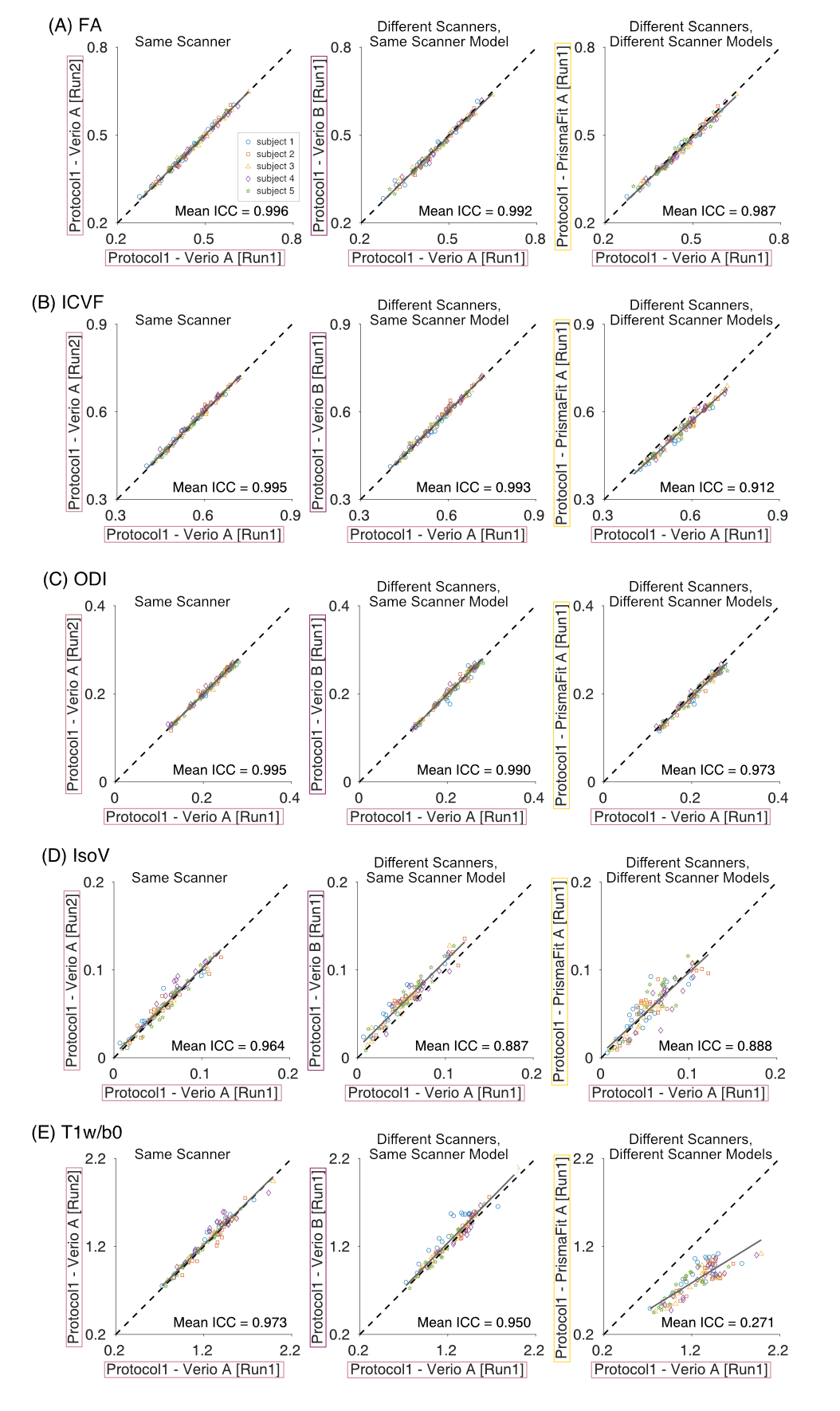

**Supplementary Figure 1. Comparison of tissue properties within a scanner, across scanners of the same model, and across different scanner models using the same acquisition protocol (protocol 1).** Each panel compares datasets acquired using the same scanner (Verio A; left panels), different scanners of the same model (Verio A and Verio B; middle panels), and scanners of different models (Verio A and PrismaFit A; right panels). Tissue-property metrics derived from tractometry analysis are shown as follows: (**A**) FA, (**B**) ICVF, (**C**) ODI, (**D**) IsoV, and (**E**) T1w/b0. All conventions are identical to those in Figure 4.

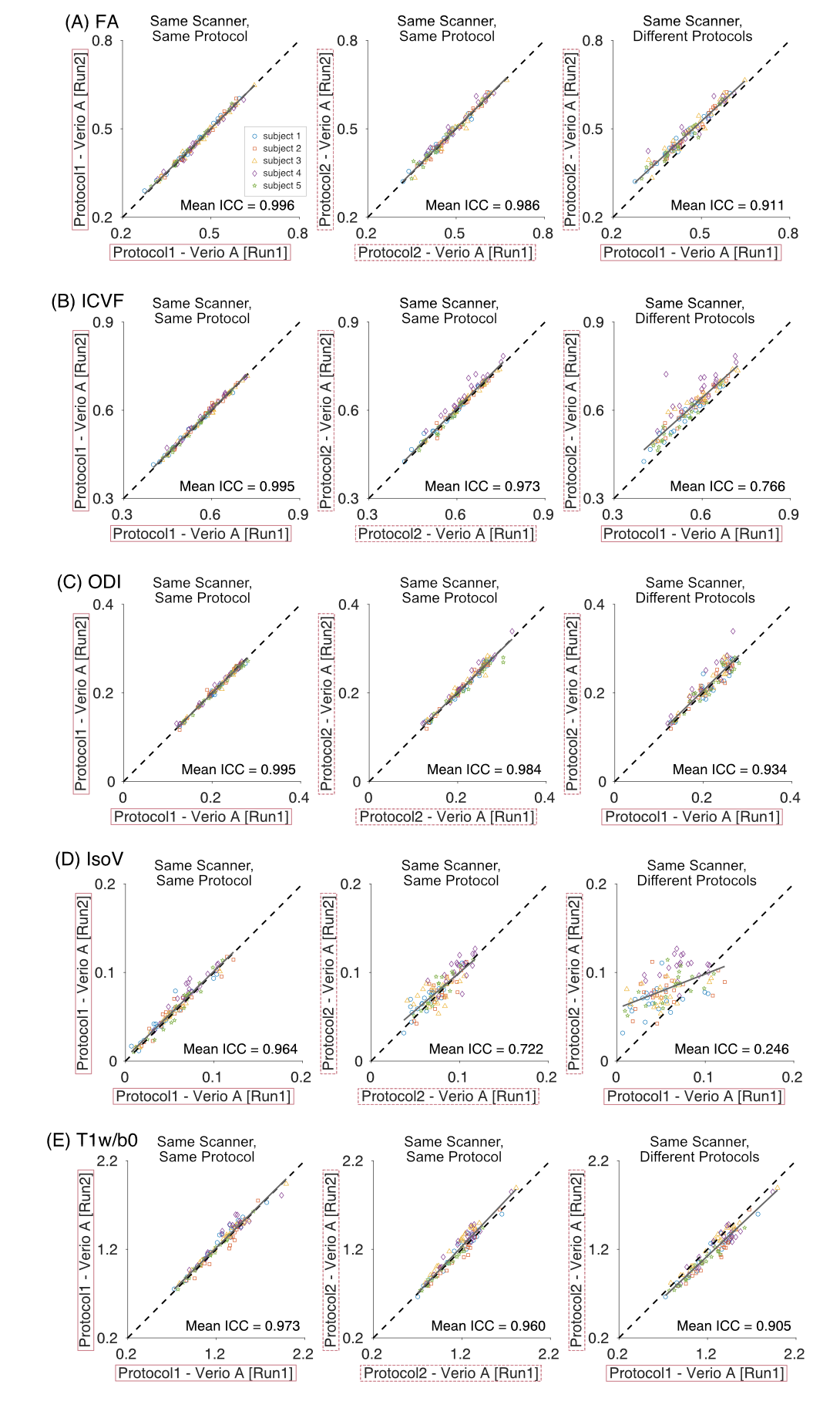

**Supplementary Figure 2. Comparison of tissue properties across datasets acquired using the same scanner with the same or different acquisition protocols.** Each panel compares datasets acquired using the same scanner and the same protocol (left panels, protocol 1; middle panels, protocol 2) and datasets acquired using the same scanner but different protocols (right panels). Tissue-property metrics derived from tractometry analysis are shown as follows: (**A**) FA, (**B**) ICVF, (**C**) ODI, (**D**) IsoV, and (**E**) T1w/b0. All conventions are identical to those in Figure 5.

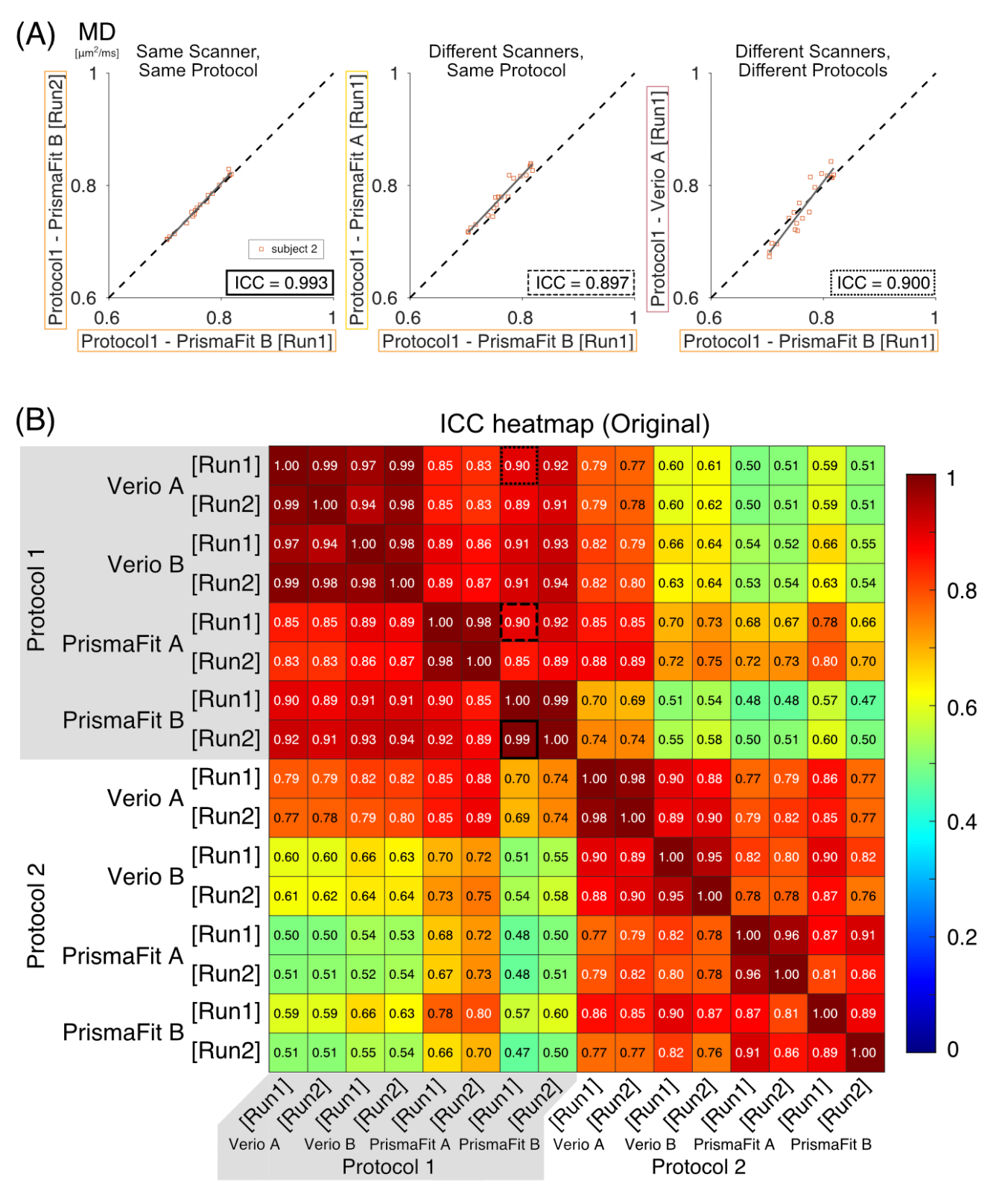

**Supplementary Figure 3. Intraclass correlation coefficient (ICC) of MD across all dataset combinations in a single subject, including PrismaFit B.** (**A**) Each panel compares MD datasets from subject 2 acquired using the same scanner (PrismaFit B; left panels), different scanners of the same model (PrismaFit B and PrismaFit A; middle panels), and scanners of different models (PrismaFit B and Verio A; right panels). All conventions are identical to those in Figure 4. (**B**) Heatmaps show ICC values for MD across all pairwise comparisons in subject 2. Rectangles highlighted with black solid and dashed lines correspond to the comparisons shown in panel A.

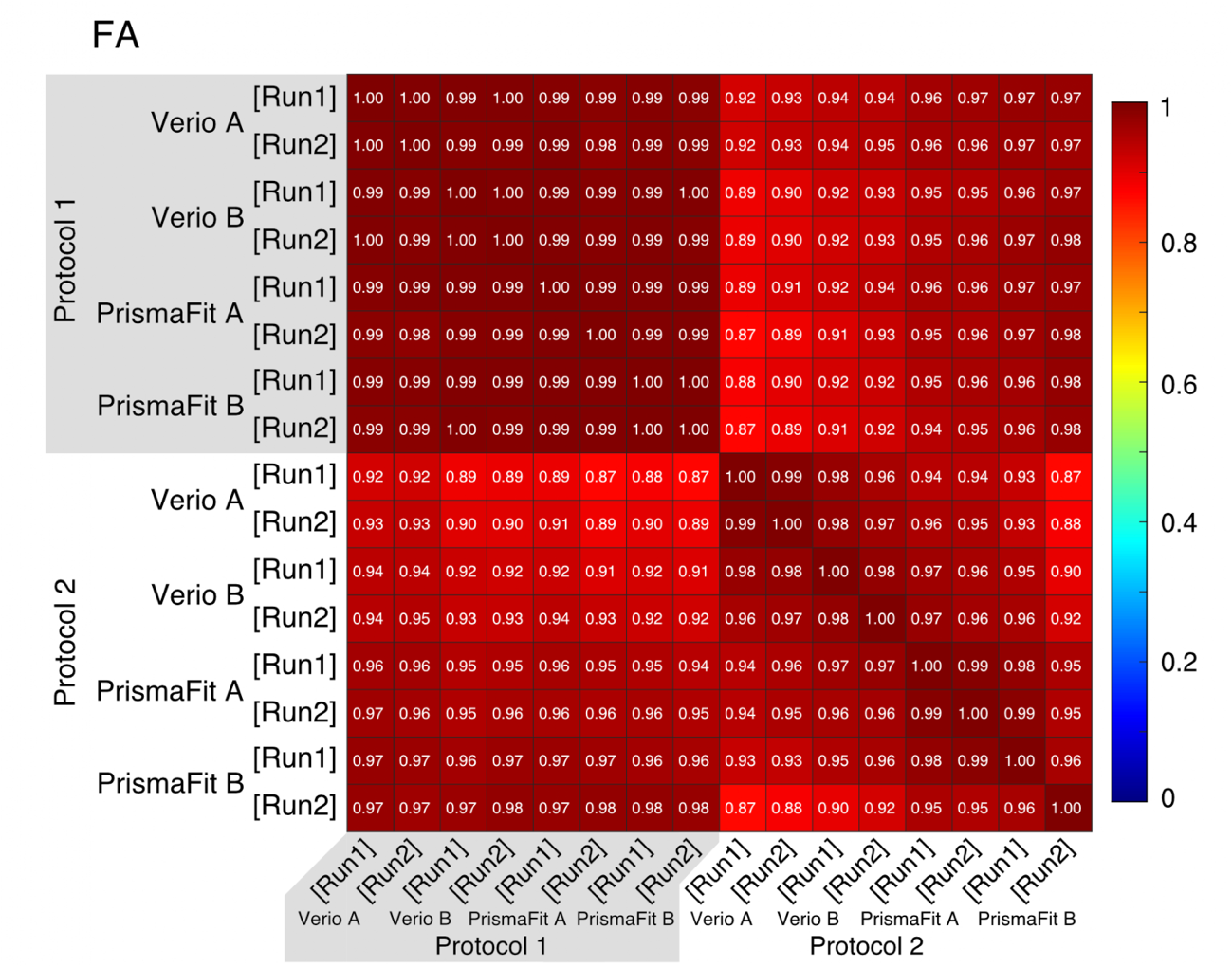

**Supplementary Figure 4. ICC of FA across all dataset combinations in a single subject, including PrismaFit B.** Heatmaps show ICC values for FA derived from tractometry analysis across all pairwise comparisons in subject 2.

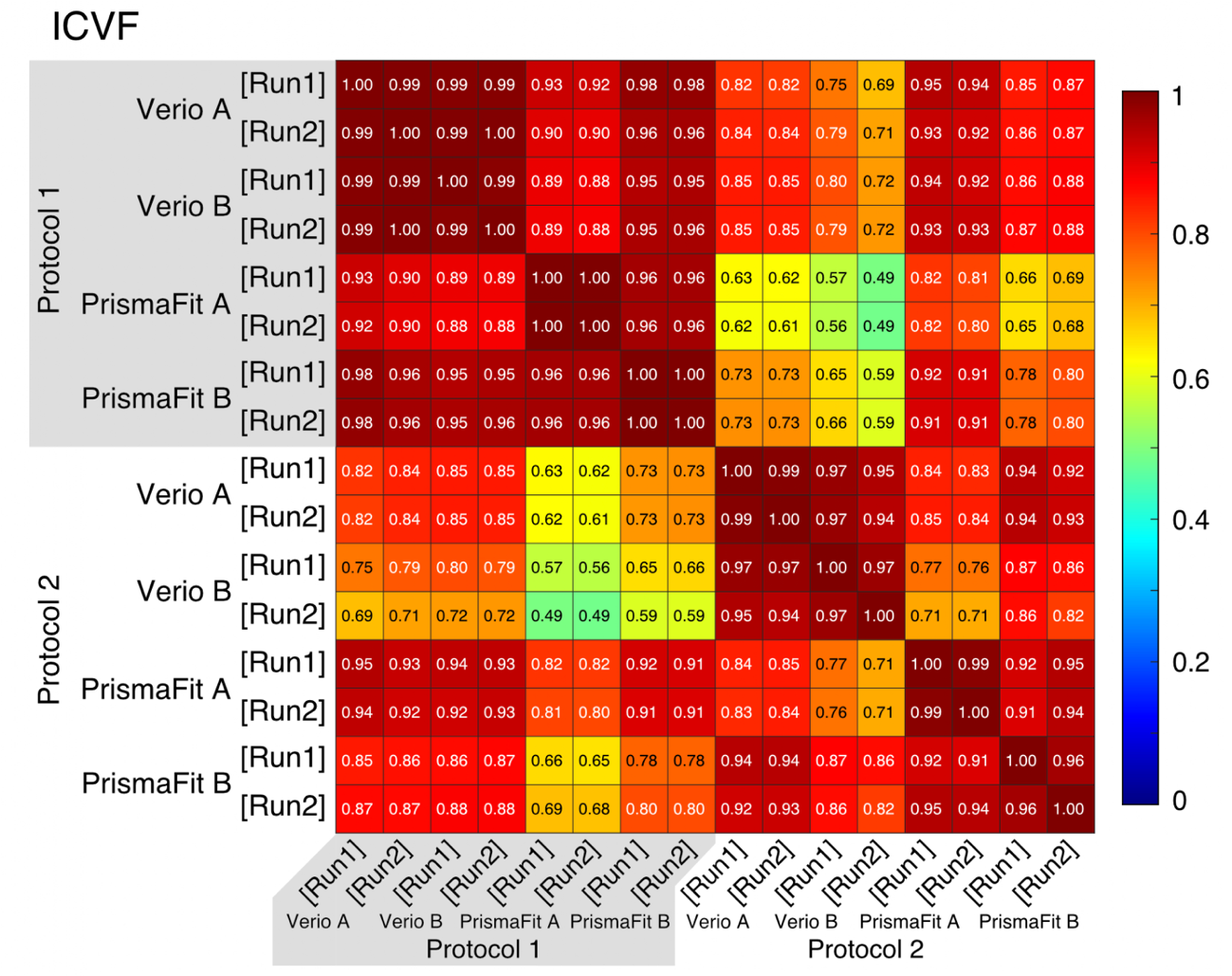

**Supplementary Figure 5. ICC of ICVF across all dataset combinations in a single subject, including PrismaFit B.** Heatmaps show ICC values for ICVF derived from tractometry analysis across all pairwise comparisons in subject 2.

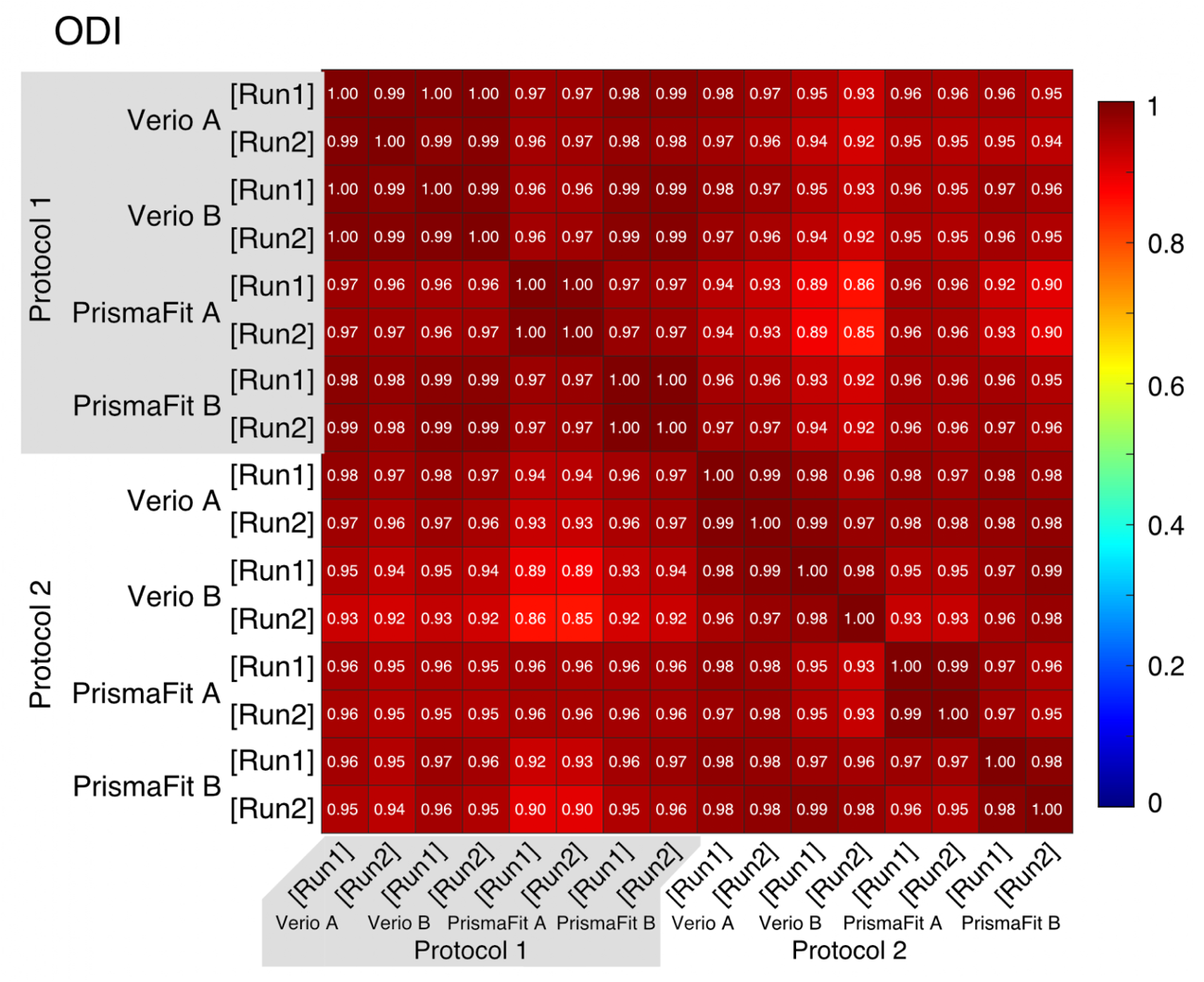

**Supplementary Figure 6. ICC of ODI across all dataset combinations in a single subject, including PrismaFit B.** Heatmaps show ICC values for ODI derived from tractometry analysis across all pairwise comparisons in subject 2.

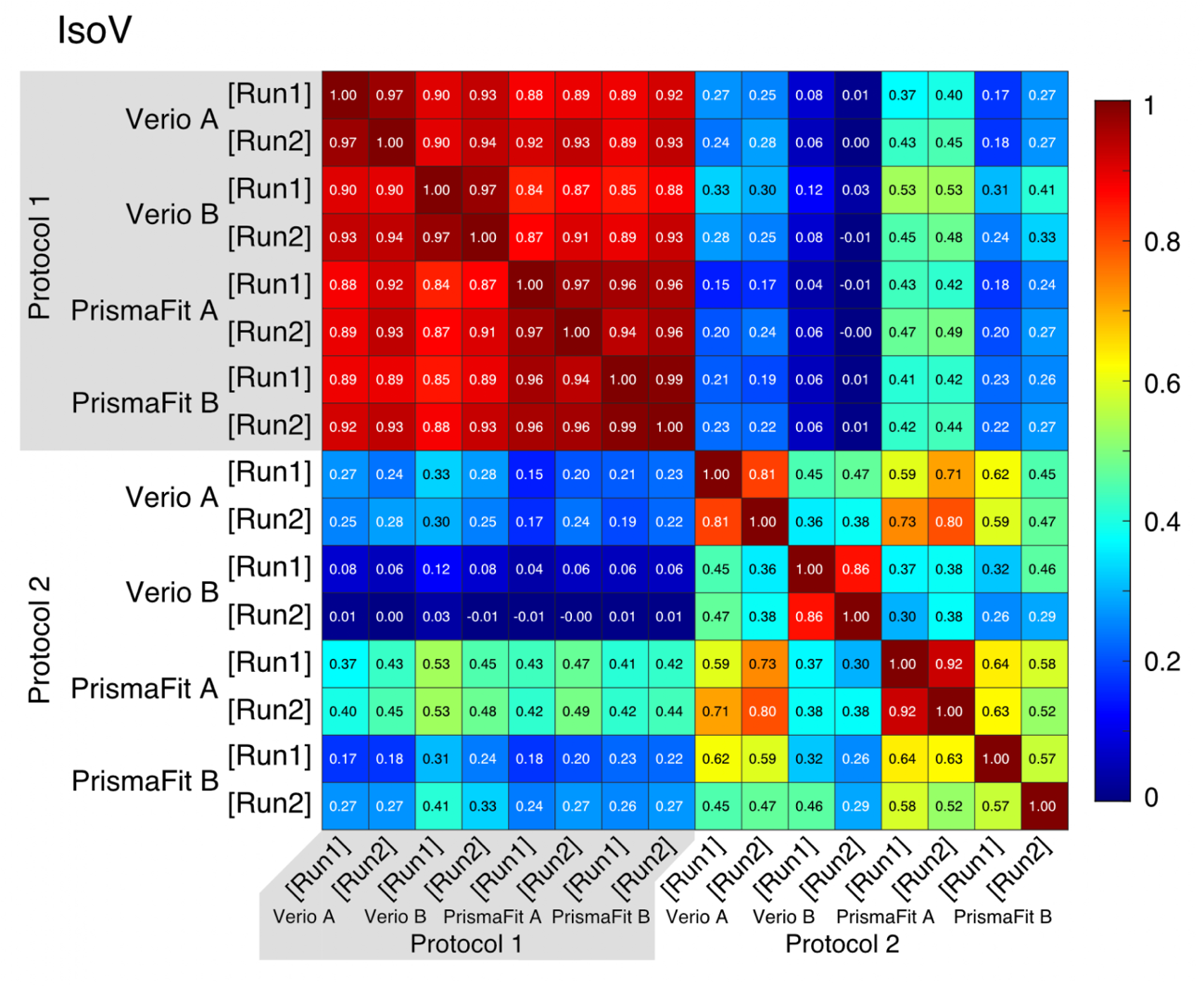

**Supplementary Figure 7. ICC of IsoV across all dataset combinations in a single subject, including PrismaFit B.** Heatmaps show ICC values for IsoV derived from tractometry analysis across all pairwise comparisons in subject 2.

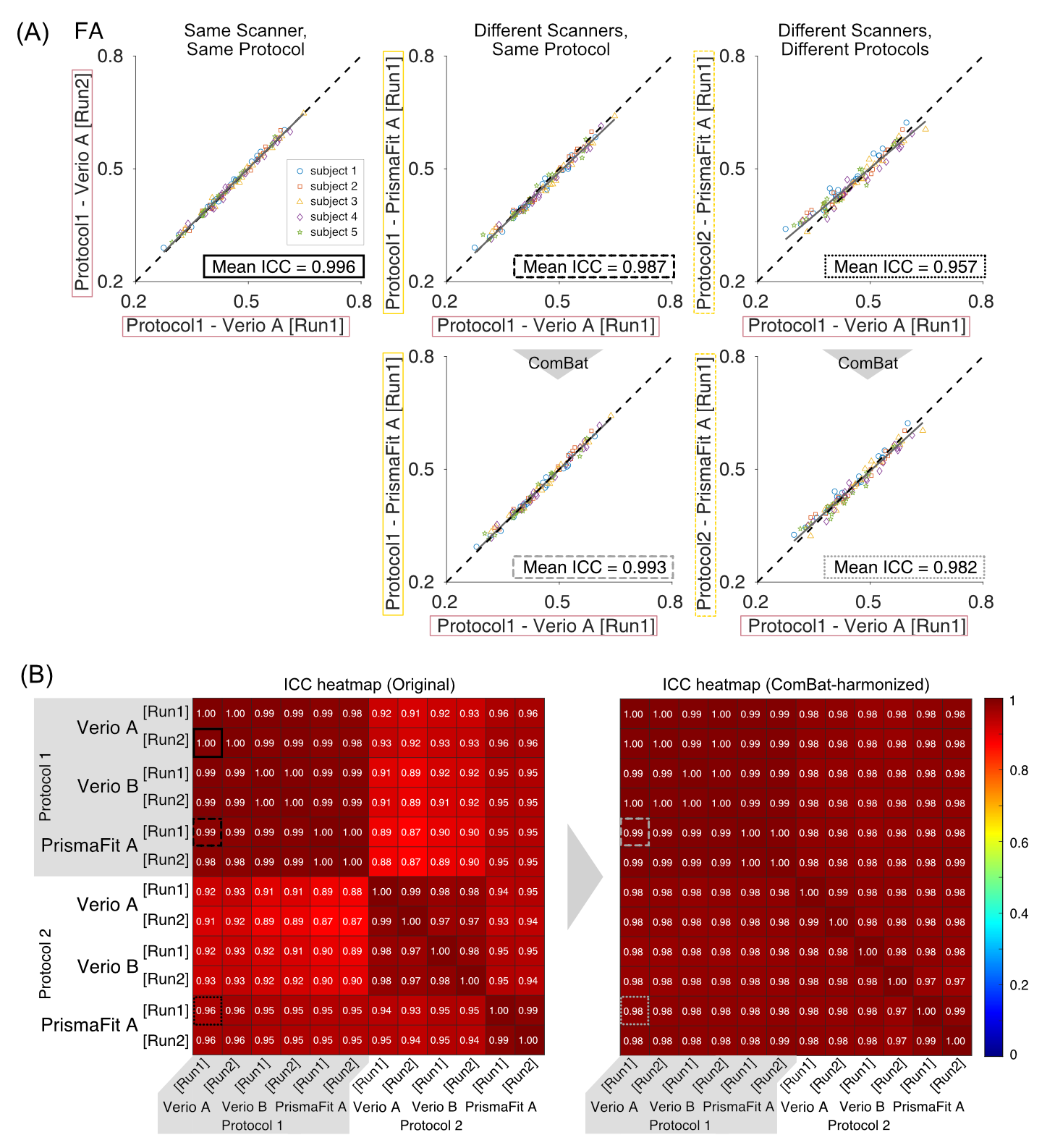

**Supplementary Figure 8. ICC of FA across all dataset combinations before and after ComBat harmonization.** All conventions are identical to those in Figure 6.

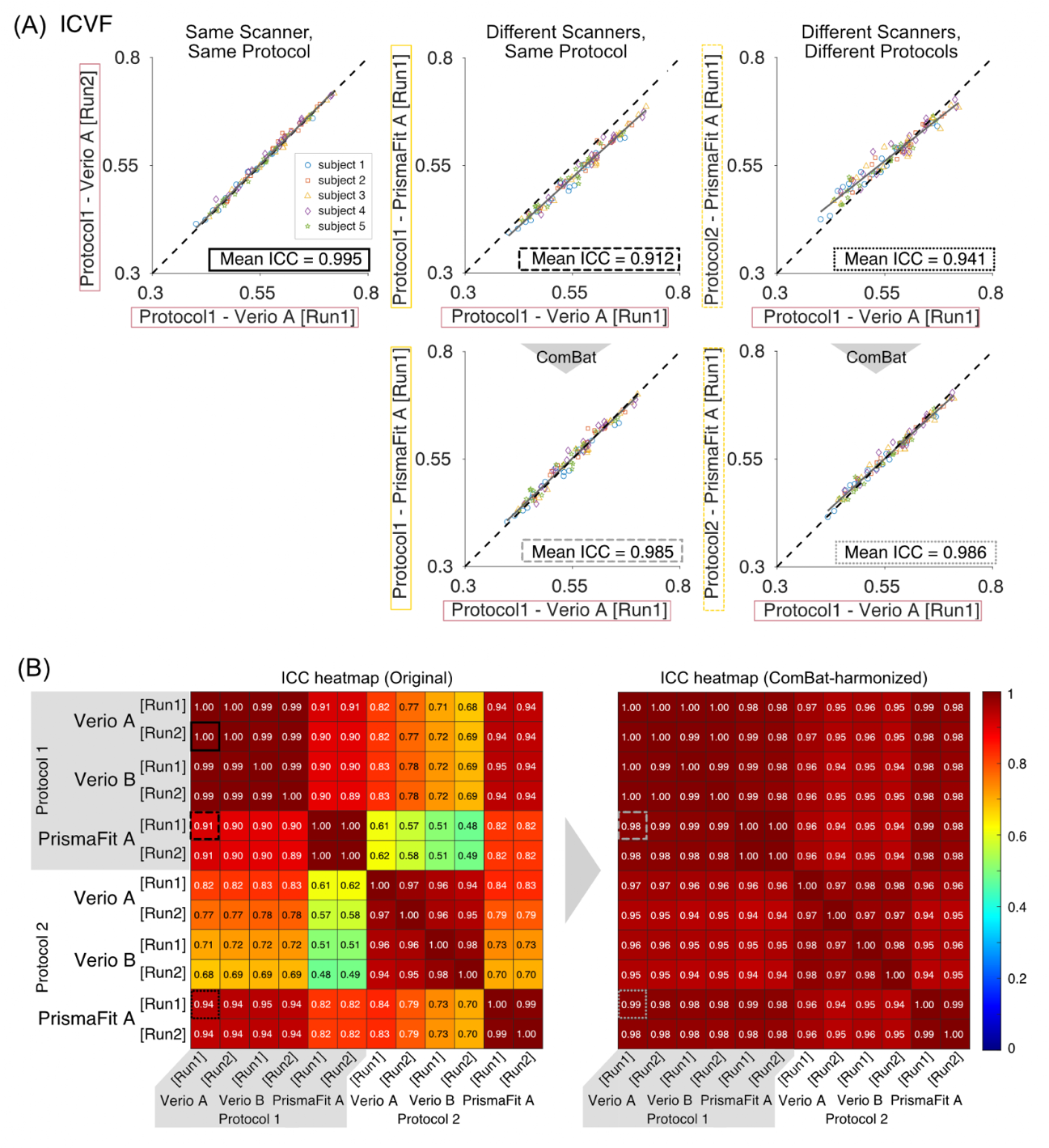
 **Supplementary Figure 9. ICC of ICVF across all dataset combinations before and after ComBat harmonization.** All conventions are identical to those in Figure 6.

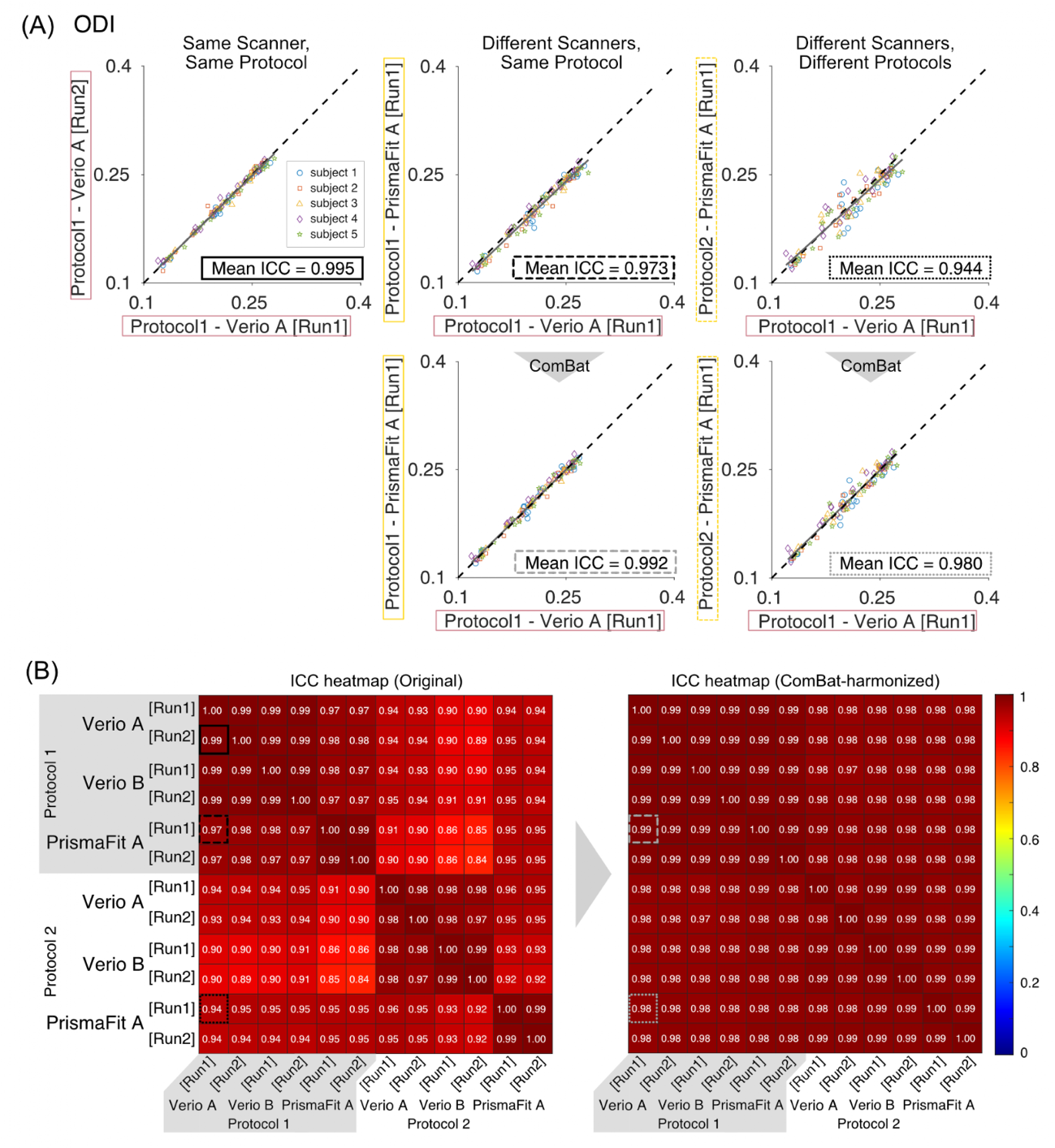

**Supplementary Figure 10. ICC of ODI across all dataset combinations before and after ComBat harmonization.** All conventions are identical to those in Figure 6.

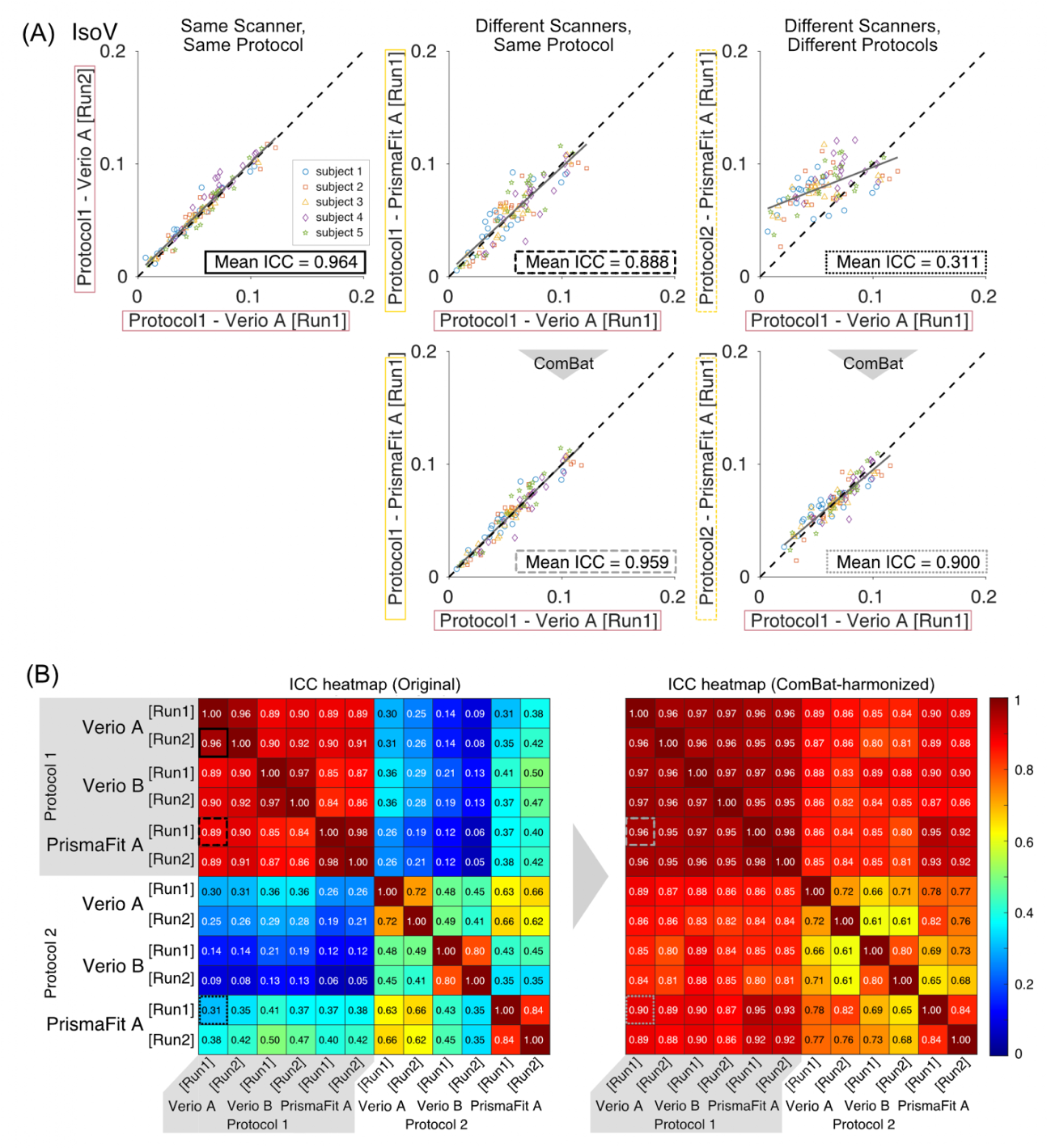

**Supplementary Figure 11. ICC of IsoV across all dataset combinations before and after ComBat harmonization.** All conventions are identical to those in Figure 6.

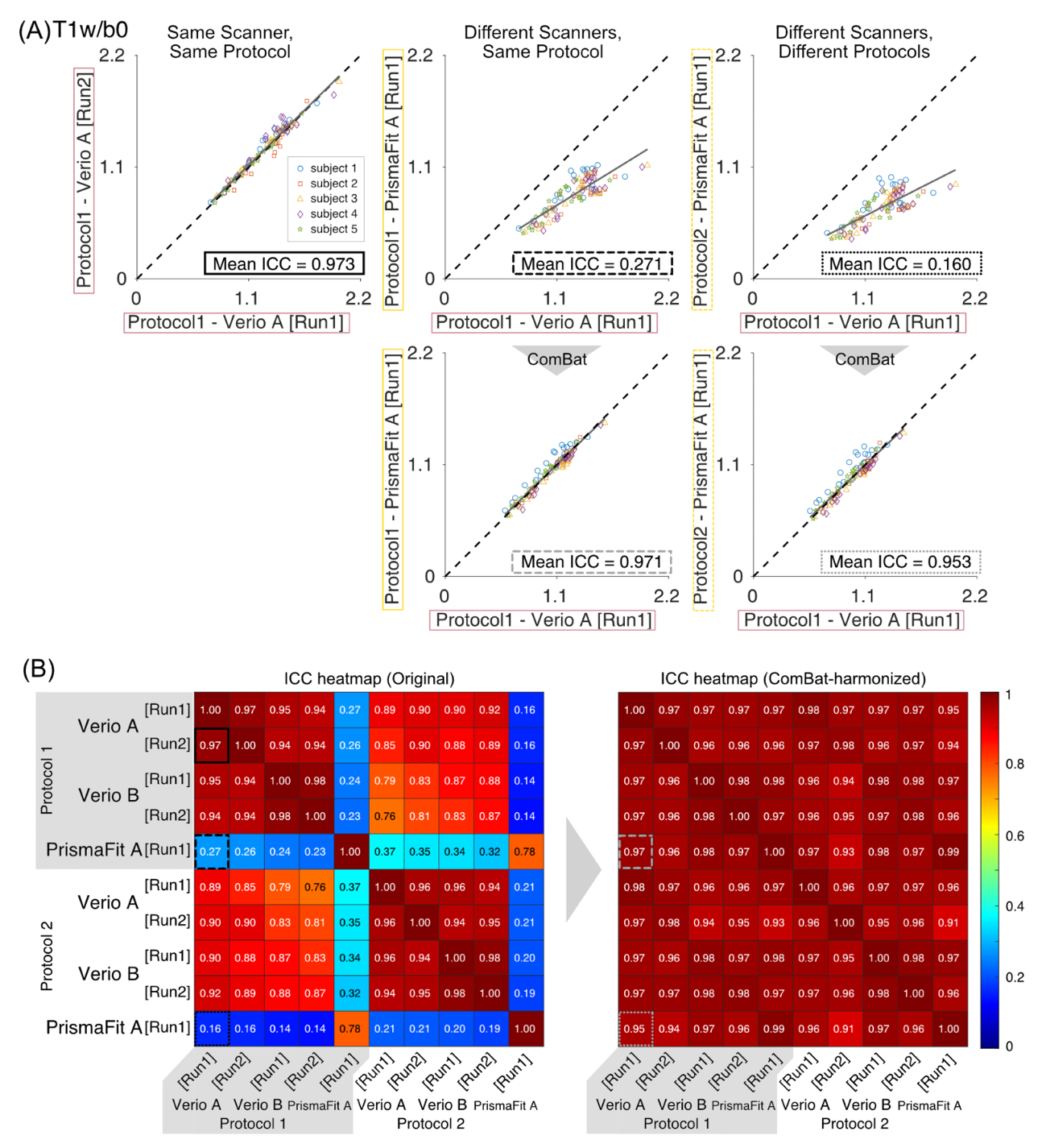

**Supplementary Figure 12. ICC of T1w/b0 across all dataset combinations before and after ComBat harmonization.** All conventions are identical to those in Figure 6.
